# Whole animal modelling reveals neuronal mechanisms of decision-making and reproduces unpredictable swimming in frog tadpoles

**DOI:** 10.1101/2021.07.13.452162

**Authors:** Andrea Ferrario, Andrey Palyanov, Stella Koutsikou, Wenchang Li, Steve Soffe, Alan Roberts, Roman Borisyuk

## Abstract

Animal behaviour is based on interaction between nervous, musculoskeletal and environmental systems. How does an animal process sensory stimuli, use it to decide whether and how to respond, and initiate the locomotor behaviour? We build the whole body computer models of a simple vertebrate with a complete chain of neural circuits and body units for sensory information processing, decision-making, generation of spiking activities, muscle innervation, body flexion, body-water interaction, and movement. Our Central Nervous System (CNS) model generates biologically-realistic spiking and reveals that sensory memory populations on two hindbrain sides compete for swimming initiation and first body flexion. Biomechanical 3-dimensional “*Virtual Tadpo*le” (VT) model is constructed to evaluate if motor outputs of *CNS model* can produce swimming-like movements in a volume of “water”. We find that whole animal modelling generates reliable and realistic swimming. The combination of *CNS* and *VT models* opens a new perspective for experiments with immobilised tadpoles.

## INTRODUCTION

Animal behaviour is based on interaction between the nervous system, musculoskeletal body units, and environment. How does an animal process sensory stimuli, use it to decide whether and how to respond, and then initiate and carry through the locomotor response? In a wide range of animals there has been progress in understanding the networks generating rhythmic locomotion (Kiehn, 2016) but the circuits that initiate it, are not well understood. Locomotor behaviour like swimming requires: brainstem/hindbrain integration of sensory stimuli; sensory memory; a decision to respond; a choice of direction; avoiding simultaneous activity of antagonists (Bahl and Engert, 2020; Gatto and Goulding, 2018; Goulding, 2009; Ruder and Arber, 2019). Although in many cases the role of neuronal populations in movement control has been identified (Bouvier et al., 2015; Kiehn, 2016), a precise description of how the sensory information is transformed to motor decision is unclear. In deciding whether to start to walk, fly or swim, all animals evaluate their sensory situation and initiate appropriate rhythmic activity. We are now in a unique position to show how this is done.

This paper describes computer modelling of a simple vertebrate with a complete chain of neuronal circuits from sensors through decision to locomotion. The model includes the spinal cord to generate rhythmic activity, adds the brainstem and sensory pathways upstream to make decisions as well as the biomechanical body and muscles downstream to reproduce swimming-like movements. The model of hatchling *Xenopus* tadpole is built at a unique level of detail compared with existing models (Abbott et al., 2016).

Frog tadpoles, one of the most numerous vertebrate animals on earth, are simple, vulnerable animals that rely on swimming to escape from predators. The simplicity of the hatchling *Xenopus* tadpole and its nervous system presents a rare opportunity for experimental and computational studies (Roberts et al., 2014; Roberts et al., 2010b) where detailed information is available on behaviour, sensory systems, the central nervous system (*CNS*) (Roberts et al., 2019), body components and their properties as well as body-water interaction (Hoff and Wassersug, 2000).

We ask how neurons in the brainstem respond to sensory signals to generate and direct locomotor behaviour (Kiehn, 2016). What role does sensory memory play in the decision to move (Goulding, 2009; Wang, 2012)? How is unpredictability of movement initiation and direction produced (Bouvier et al., 2015; Li et al., 2015)? What are the basic theoretical principles guiding these responses (Merel et al., 2019)? These key questions are addressed using computer modelling of a simple vertebrate.

The *CNS model* includes two skin touch sensory pathways to initiate swimming, an inhibitory head skin pressure pathway to stop it, and hindbrain populations of sensory memory neurons (*exIN*s) which prolong signals from sensory pathways and turn on the swimming Central Pattern Generator (CPG). We focus on the critical role of *exIN*s which provide a basic sensory memory of the brief stimulus (Wang, 2012). CNS model simulations reveal that the neuronal mechanism of decision is based on an accumulation of competing excitations to threshold (Brody and Hanks, 2016; Carpenter and Williams, 1995; Gold and Shadlen, 2007; Noorani and Carpenter, 2016) on two body sides. In particular, simulations explain experimental evidence (Roberts et al., 2019) on long and variable delays between stimulation and the start of swimming as well as the side of the first body flexion. Introducing unpredictability in the side of first flexion may reduce predation because the direction of tadpole movement becomes unpredictable for a hunting predator (Li et al., 2014a).

We evaluate whether the motor output from the *CNS model* can produce swimming movements by building a novel 3D biomechanical computer *Virtual Tadpo*le (*VT*) model of tadpole body. The *VT model* is based on detailed physical and anatomical measurements of the shapes and mass distribution of organs like the notochord, muscles and belly in real tadpoles. We place the reconstructed virtual body in “water” and feed patterns of motoneuron spiking from the *CNS model* to the muscle segments of *VT model* to drive “virtual” swimming movements. Supplementary videos demonstrate that the simulations of *VT model* produce realistic swimming behaviour.

The combination of *CNS* and *VT models* opens a new perspective for experimental studies. Being realistic and detailed, the *CNS model* can be fitted to match experimental recordings from immobilised tadpoles. Motoneuron spiking patterns generated by the fitted *CNS model* can activate muscle segments of *VT model* to test movement in water. This novel approach leads to new theoretical insights for experimental testing.

Here we provide an introductory background on **tadpole swimming and its neuronal control**. When a hatchling *Xenopus* tadpole is touched, it flexes unpredictably to the left or right and then swims away even if the mid- and fore-brains are disconnected (Fig. 1a,b,d). Swimming stops when the head bumps into solid support and attaches using cement gland mucus (Perrins et al., 2002). Tadpoles swim belly down because dense ventral belly cells act as ballast (Roberts et al., 2000). Segmented muscles lie on each side of the spinal cord and a notochord acts as a longitudinal skeletal rod.

**Figure 1:**
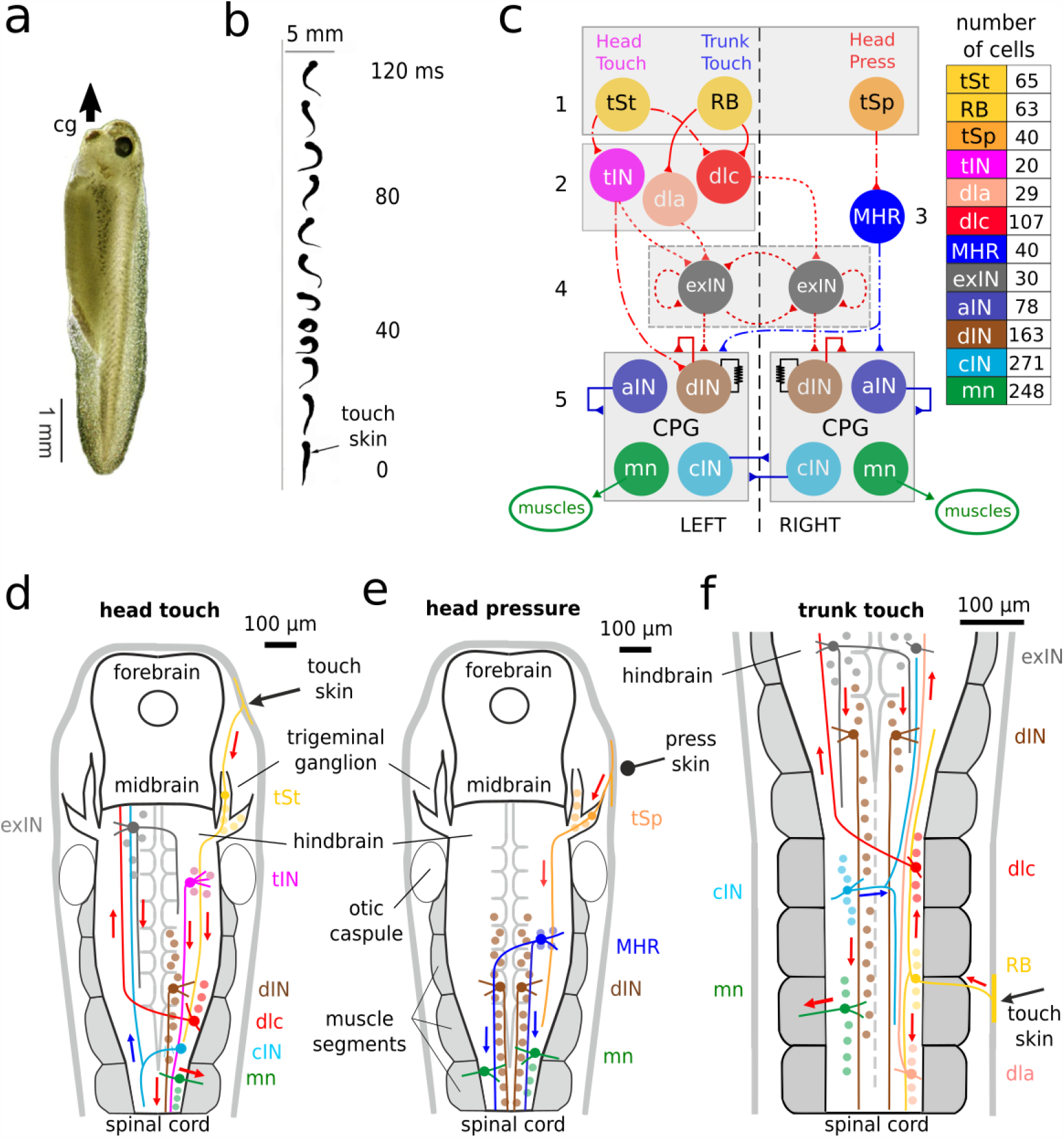
Tadpole, swimming and neuronal pathways. **a**. Tadpole hanging from mucus (arrow) secreted by cement gland (cg). **b**. If touched (arrow), it flexes to one side and then swims off. **c**. Functional diagram of *CNS model* network with 12 neuron types forming 5 layers: 1) skin sensory; 2) sensory projection; 3) inhibitory reticulospinal; 4) sensory memory; 5) central pattern generator. The list shows colour coding and population of each neuron type on one side. The total number of neurons in CNS model is 2,308 (half on each side) including 788 sensory pathway and 1,520 CPG neurons. Connections shown by solid lines have been established by “developmental” modelling based on single cell recording and dye injections (Roberts et al., 2014); dash-dotted and dotted lines indicate connections prescribed by the “probabilistic” model (dashed line means that connections probabilities are based on some limited experimental evidence). Red are excitatory and blue are inhibitory connections; resistor sign shows electrical connections. Abbreviations not defined in text: Interneurons: *aIN* = ascending; *cIN* = commissural; *tIN* = trigeminal; *dla* = dorsolateral ascending; *dlc* = dorsolateral commissural; *mn* = motorneuron. **d-f**. Diagrams of the brain and spinal cord illustrating the neuronal pathways from head (**d**) or trunk touch (*f*) to start and from head pressure (**e**) to stop swimming. Only selected neurons are shown to make the signal propagation pathways clear: coloured arrows show activity propagation pathways (red = excitation; blue = inhibition). The hindbrain extends from 0 - 500 microns and the spinal cord from 500 - 3500 microns.

Detailed information on the neurons initiating and controlling swimming comes from paired whole-cell recordings in immobilised tadpoles with dye injection to reveal neuron anatomy (Roberts et al., 2010a) (overview Fig. 1c-f;. Neurons are mainly arranged in longitudinal rows, have short local dendrites and longitudinal axons projecting on the same or the opposite side. Motor nerve recordings reveal swimming activity to 1ms electrical stimuli which excite sensory neurons in the head (*tSt*) or trunk (*RB*) skin. Sensory pathway neurons then transmit brief excitation to the hindbrain on both sides of the body (Buhl et al., 2012). A slow wave of excitation builds up in electrically coupled hindbrain reticulospinal descending *dIN*s until firing threshold is reached on one side of the body. The *dIN* population on that side is recruited to fire, excites motoneurons (*mn*s) and swimming starts (Buhl et al., 2015; Koutsikou et al., 2018; Li et al., 2004a).

The hindbrain *dIN*s neurons make the decision to swim and drive the other swimming CPG neurons including motoneurons (Figs. 1 c,d,f). They are probable homologues of brainstem neurons in zebrafish larvae (Bahl and Engert, 2020). We have proposed a preliminary hypothesis that: (1) the wave of excitation in hindbrain *dIN*s comes from populations of hindbrain sensory memory, extension neurons (*exIN*s) on each side of the body which are excited following skin stimulation (James, 2009) (level 4 in Fig. 1c); (2) The *exIN*s then fire for about 1s as a result of proposed mutual re-excitation (Koutsikou et al., 2018). Variation in the firing of the *exIN*s on each side of the body leads to unpredictability in the side where *dIN* firing threshold will be reached and the first flexion of swimming will occur. Sustained swimming then depends on self-re-excitation within the *dIN* populations on each side of the body and on rebound firing following mid-cycle inhibition from the reciprocal inhibitory *cIN*s on the opposite side (Roberts et al., 2014).

As swimming continues, the cycle period lengthens and stopping can occur spontaneously (Dale, 2002) or when pressure on the head skin excites trigeminal sensory neurons (*tSp*s). These excite mid-hindbrain GABAergic inhibitory reticulospinal neurons (*MHR*s) which project caudally on both sides of the body to inhibit CPG neurons and stop swimming (Perrins et al., 2002) (Figs. 1c, e).

## RESULTS

### HOW CNS *MODEL* GENERATES SWIMMING BEHAVIOUR

The new *CNS model* is a network of 2,308 spiking, Hodgkin-Huxley type neurons of 12 different types (Fig. 1 c-f). This model can generate a complete sequence of swimming behaviour controlled by skin sensory inputs (Fig. 2) which is very similar to real tadpole behaviour in response to natural stimuli. It produces motoneuron activity patterns for swimming initiated by “touch” to the head or trunk skin which evokes brief firing in the sensory pathways (Fig. 2b, d). Swimming can be terminated by head skin “pressure” when sensory and inhibitory reticulospinal neurons (*MHR*s) fire repetitively (Fig. 2c). As swimming continues, cycle period lengthens but can be accelerated by a trunk skin stimulus (Dale, 2002; Li and Moult, 2012; Roberts et al., 2014) (Fig. 2a, bottom). The initial shorter cycle periods can be explained by a higher depolarisation of rostral *dIN*s due to excitation from active *exIN*s lasting for about 1s. This depolarisation leads to post-inhibitory rebound firing after *cIN* spiking. Swimming can also stop spontaneously after post-inhibitory rebound failure.

**Figure 2:**
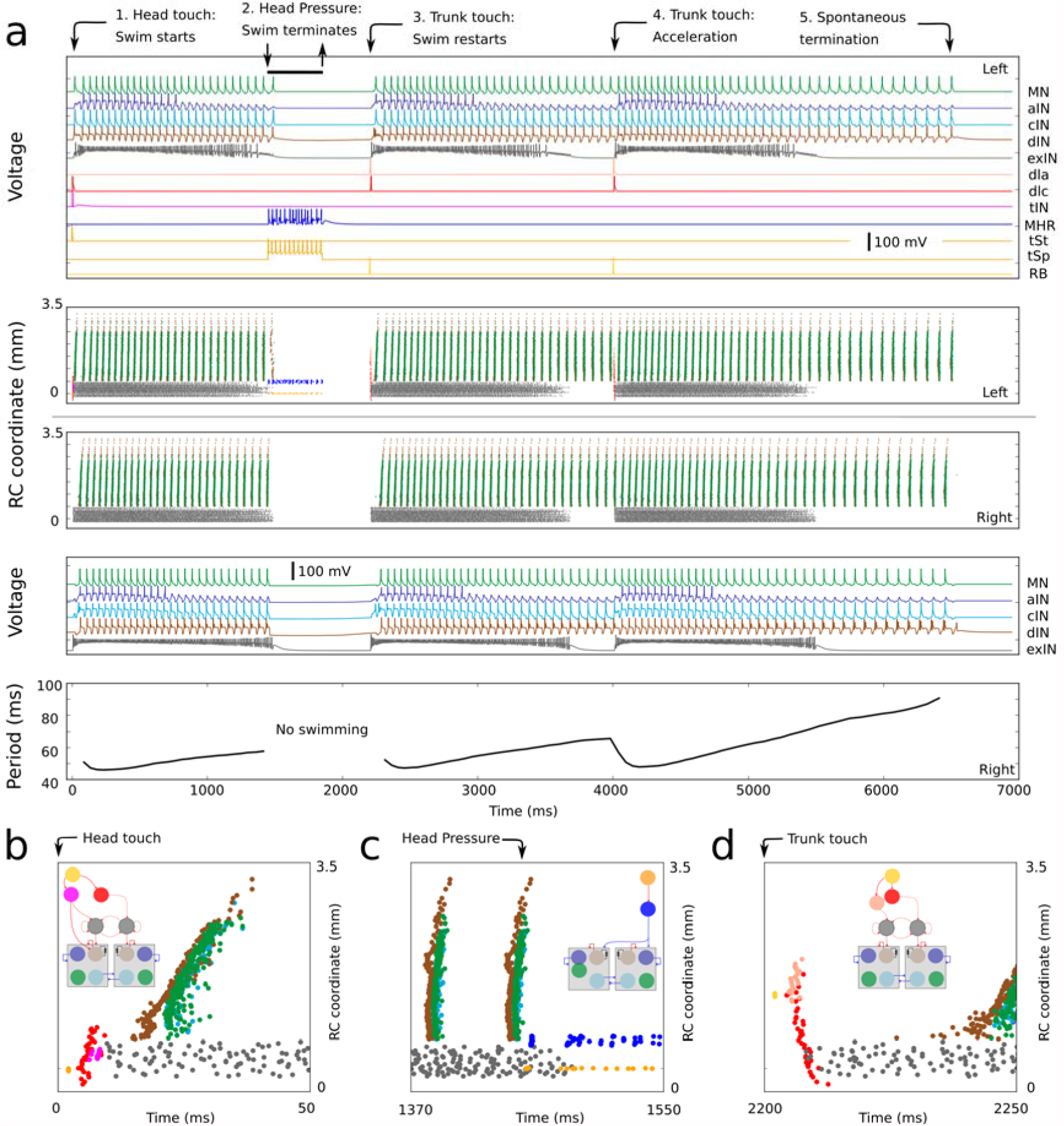
Overview of CNS *model* responses to a sequence of sensory signals. **a**. The “Voltage” panels show membrane potentials of selected active neurons on the left and right sides respectively. The two central panels show spike times (coloured dots, zoomed views in *b-d*) of all active neurons, the vertical coordinate is the rostro-caudal position of the spiking neuron. All current pulse sensory stimuli are applied to the left side of the body as indicated by black arrows at the top. 1) Head skin touch initiates swimming from the rest (stimulus of 0.3 nA for 5ms excites 13 *tSt*s at time zero); 2) Head skin pressure stops swimming (stimulus 0.3nA for 30ms excites 10 *tSp*s at time 1.3s; 3) Trunk skin touch initiates swimming (stimulus of 0.3nA, for 5ms excites 2 *RB*s at time 2s) ; 4) Trunk skin touch during swimming leads to acceleration (stimulus of 0.3nA, for 5ms excites 2 *RB*s at time 4s); 5) Swimming slows and stops spontaneously. **b-d**. Zoomed in view of spike times for neurons on the left side responding to each type of skin stimulation. Inserts show small diagrams of each sensory pathway (see Fig. 1c).

Our previous model of the tadpole network (Roberts et al., 2014) was based on extensive anatomical and electrophysiological data from experimental studies (Ritson and Li, 2019; Roberts et al., 2010a). It included CPG neurons and generated swimming type spiking activities triggered by trunk skin stimulation. Biologically realistic pair-wise connectivity was modelled using a “developmental method” (Borisyuk et al., 2014) and was then mapped onto a functional model. Swimming initiation was simplified to start after a short delay and stopping was not included. The *CNS model* is enlarged to include decision making for swimming initiation (on stimulated or opposite side) with a long and variable time delay. Thus, the new model generates the appropriate spiking dynamics and reproduces corresponding movements in response to any sequence of sensory inputs.

### *CNS MODEL* DESIGN

Here we extend our previous swimming model (Roberts et al., 2014) by adding 2 sensory, 2 sensory pathway and 1 reticulospinal neuronal populations (*tSt, tSp, tIN, MHR, exIN*) responsible for starting and stopping swimming (Fig. 1c). The connectivity of these neurons was established using the probabilistic method (Ferrario et al., 2018) as well as a novel technique of anatomical and functional similarities between pairs of new and existing populations. Doing that we hypothesise that for two similar pairs of populations, the connection probabilities in each pair are inferred from the same probabilistic distribution. Details of two new algorithms for finding connection probabilities are in Methods.

Our previous swimming model (Roberts et al., 2014) used extensive anatomical data to partially reconstruct the neuronal connectome and physiology of the sensory pathway and CNS neurons (*RB, dla,dlc, dIN, cIN, aIN, mn*). Further work (Ferrario et al., 2018) used this connectivity to extract the probability of each pairwise connection in the connectome (probabilistic model).

Crucially, we included new populations of unidentified hindbrain neurons (*exIN*s) which have been proposed to explain the slow build-up of excitation to threshold leading to swimming in response to trunk or head skin touch (Koutsikou et al., 2018). The connections between *exIN*s and other populations are assigned in one of three ways: (1) To calculate *dla*/*dlc* to *exIN*s connection probabilities we use recordings from irregular spiking midbrain neurons (Methods) assuming their properties are similar to the *exIN*s. (2) We explore the probability space to obtain long lasting irregular *exIN* firing and reliable initiation of swimming. (3) To optimise the probability and strength of *exIN* to *hdIN* connections we use under swimming threshold stimulation and compare the response of hindbrain *dIN*s with intracellular recordings (Koutsikou et al., 2018).

**The physiological model** of each single neuron type follows the Hodgkin-Huxley formalism, with parameters that fit the electrophysiological properties of each cell type (details in Methods). Models of trunk touch sensory pathway and CPG neurons are the same as in our previous publications (Borisyuk et al., 2014; Ferrario et al., 2018; Sautois et al., 2007) except for *dIN*s. The model of *dINs* is simplified from a multi-compartmental model (Hull et al., 2015) with one soma/dendrite compartment. For the head touch sensory pathway neurons *tSt* and *tSp* we adjust the *RB* model (Sautois et al., 2007) and parameters of *MHR* and *tIN* models are selected to match the input resistance, threshold current and current-frequency curve with experiments. The *exIN* model has two compartments: soma/dendrite and axon (Koutsikou et al., 2018).

**Pairwise connectome of CNS model** is based on connection probabilities between all neurons: *P* (*i, j*)is the probability of directed connection from neuron *i* to *j,i* = 1,2,,…,*N ;j* = 1,2, …, *N, N* is the total number of neurons. By sampling from these probabilities we generate a neuron-to-neuron connectome which is represented by the binary adjacency matrix *A*(*i,j*), (*i* = 1,2, …, *N*; *j* =1,2, …, *N*),where *A* (*i, j*) = 1 means a directed connection from neuron *i* to *j*. Fig. 3a shows an example connectome with ∼ 128K pair-wise connections. Remarkably, when different connectomes were projected to the physiological model of spiking neurons, all of them generated functional behaviour that correspond to the ones found in experiments and described below.

**Figure 3:**
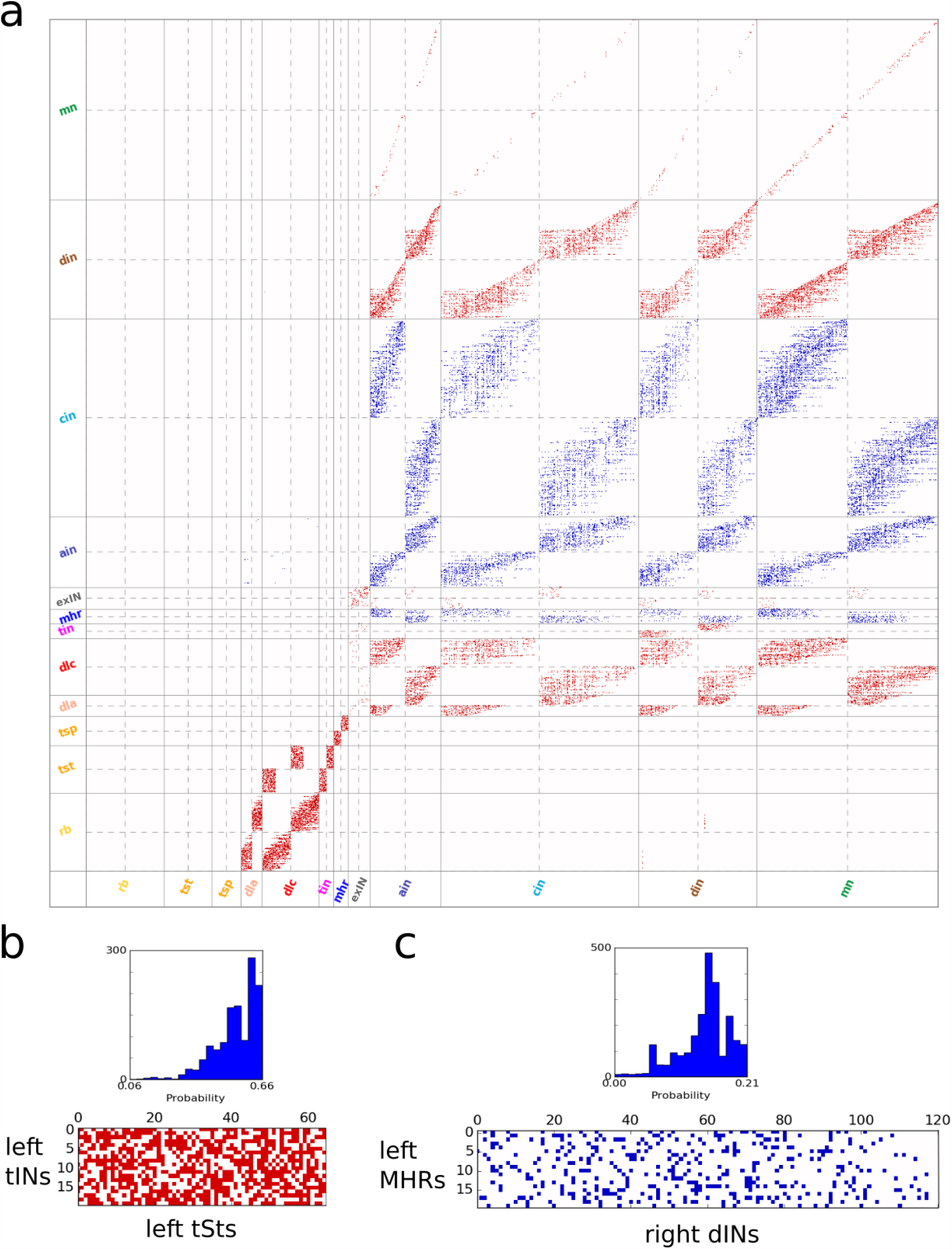
Connectivity of *CNS model*. **a**. Visualisation of the adjacency matrix (connectome), where red are excitatory and blue are inhibitory connections. Rows and columns correspond to pre- and post-synaptic neurons, respectively. There are 12 types of neuronal populations in the *CNS model* and they are separated by solid grey lines. Dashed lines separate the matrix into symmetrical sub-blocks. Within each sub-block vertical and horizontal dotted lines separate the left body side (top rows and left columns) from the right body side (bottom rows and right columns). In each sub-block neurons are ordered according to increasing rostro-caudal position. The matrix describes 128,958 pair-wise connections in one *CNS model* (in 100 models, the mean of total connection number: 128,845.8; s. d. 1,715.4). **b-c**. Examples of connection probabilities distributions (histograms) and zoomed extraction (lower panels) of excitatory (red) and inhibitory (blue) connections for *tSt->tIN* and *MHR->dIN*, respectively. **(b)** Probabilities *p*_*k*_, (*k* = 1,.., *m*) for *tSt->tIN* connections are in the range [0.06, 0.66]. Note: this sub-matrix has been extracted from the adjacency matrix *A* (*i, j*) and transposed: here columns and rows correspond to pre- and post-synaptic neurons, respectively. The number of connections *m* = 691 and the mean and standard deviation: 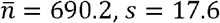 (formula (1), the mean of non-zero probabilities 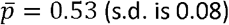 (s.d. is 0.08) - about half of possible connections. **(c)** For *MHR->dIN* connections the probabilities are in the range [0.001, 0.21]. The number of connections *m*= 338, the mean and standard deviation: 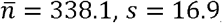, the mean of non-zero probabilities 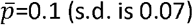 – sparse connections.

The estimate the average number of connections in the adjacency matrix from the population with *m*_1_ neurons to the population with *m*_2_ neurons we consider the Poisson binomial distribution and calculate the mean 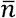 and the standard deviation *s*:

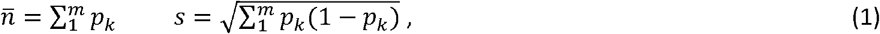

where *p*_*k*_, (*k* = 1,2, …, *m*) are probabilities of directed connections from one population to another, *m*= *m*_1_ *\*m*_2_.

Fig. 3b,c illustrates the way to derive connections from probabilities. Fig. 3b (top) shows the distribution (histogram) of connection probabilities from sensory *tSt* to *tIN* neuronal populations that is skewed towards higher probabilities. Fig. 3b (bottom) shows zoomed connections where the number of connections is in a good agreement with formulas (1). To characterise sparseness of these connections in comparison with the fully connected network we calculate the average probability 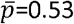 and conclude that about half of possible connections is present.

Similarly, Fig. 3c shows the histogram of connection probabilities (top) from mid-hindbrain reticulospinal *MHR*s to *dIN*s that is slightly skewed towards higher values and zoomed connections extracted from the adjacency matrix (bottom). The average probability 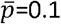 shows that connections (bottom) are rather sparse.

### THE ROLE OF SENSORY MEMORY NEURONS IN SWIM STARTS

The alternating firing of reticulospinal *hdIN*s on the two sides of the body is what drives swimming (Koutsikou et al., 2018; Soffe et al., 2009). In response to weak trunk skin stimuli, a ramp of excitation is recorded in single *hdIN*s (Koutsikou et al., 2018) (Fig 4a, b). This may lead to the start of swimming unpredictably on the stimulated or unstimulated side at variable delays after the stimulus but does not always initiate swimming. Therefore the stimulation per se does not define the start or the sidedness of the first motor reactions. Furthermore, *hdIN*s and *mn*s on both sides occasionally produce a few cycles of synchronous left and right firing (synchrony) preceding swimming alternation. Such synchrony has been studied (Ferrario et al., 2018; Li et al., 2014b) but currently appears to have no behavioural function.

**Figure 4:**
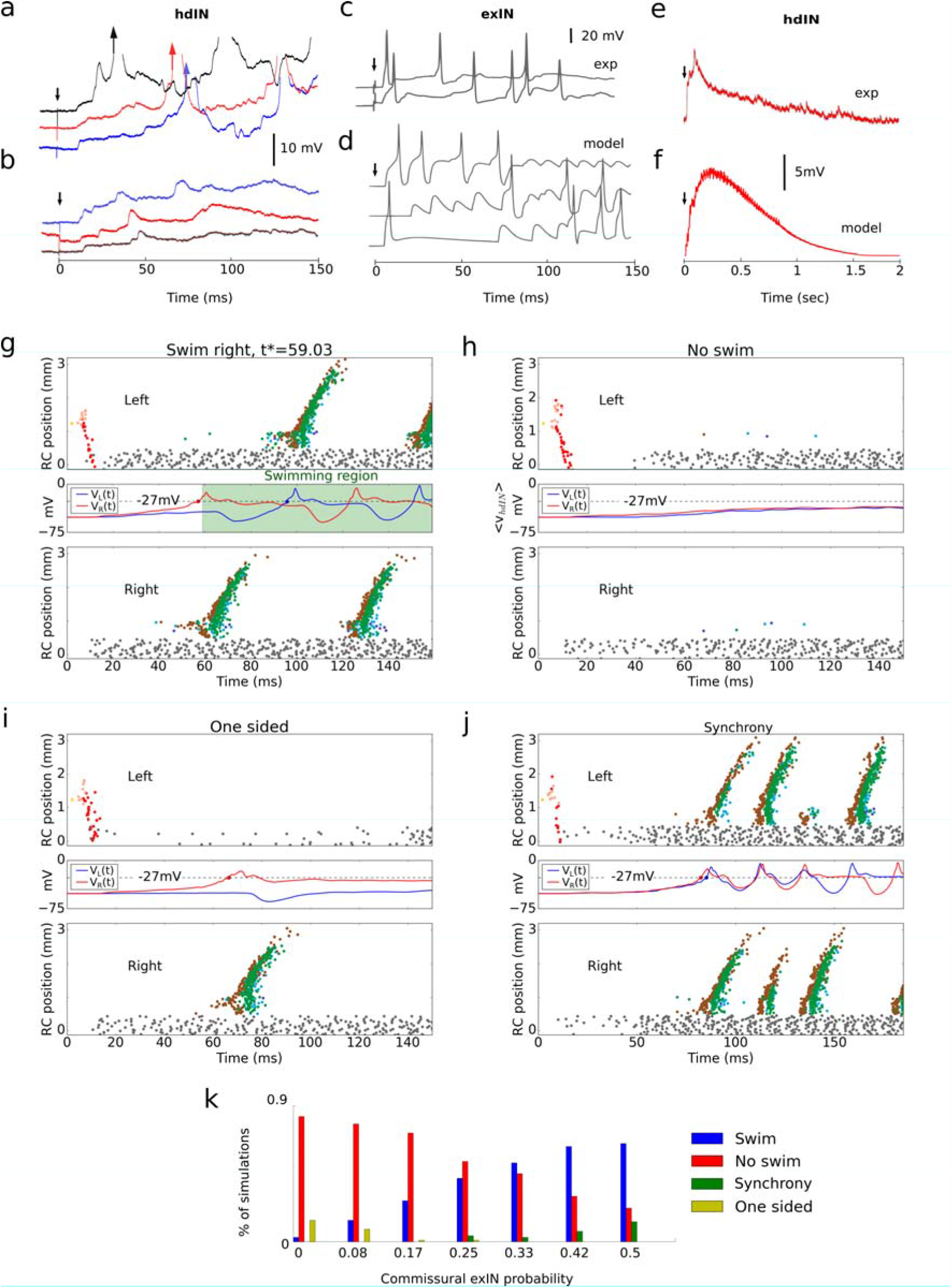
Activity of *exINs* and neuronal mechanism of swimming initiation. **a-b**. Six *hdIN* recordings of responses to head skin stimulation (black arrow) on the unstimulated side (from Fig. 6 (C-D) (Koutsikou et al., 2018)). **(a)** Excitation ramps to threshold and firing leads to swimming (arrow shows first spike). **(b)** Excitation ramp does not reach threshold. **c**. Recordings from model *exIN*s to a trunk stimulation (at arrow). **d**. Two recordings of a possible hindbrain sensory processing neuron response to a trunk skin stimulation (at arrow; from Fig 3(g) (Roberts et al., 2019)). **e-f**. The averaged ramp of 5 randomly selected model *hdIN*s **(e)** and 5 experimentally (exp) recorded *hdIN*s **(f)** (from Fig. 6F (Koutsikou et al., 2018)) to a subthreshold trunk skin stimulation (at arrow). In the model, the ramp decays significantly by 1.5 s due to synaptic depression of *exIN*s interconnections. The strength of *exIN*->*hdIN* connection is selected to match the average *hdIN* potential. **g-j**. *CNS model* responses to trunk stimulation: swimming **(g)**, no response **(h)**; one sided activity **(i)**; synchrony **(j)**. Top and bottom subpanels show spike times of active neurons. Central panel shows the averaged voltage dynamics of *hdIN*s on left *V*_*L*_ (*t*), blue) and right *V*_*R*_ (*t*), red) sides. Coloured dots indicate crossing the threshold (dotted line). In the case of swimming *(g)*, green area corresponds to swimming activity and initiation time *t*^*^ is the mean of *hdIN* spike times on right. **k**. Bar chart shows distribution of four responses for different connection probabilities between *exIN*s on opposite sides. For each probability we run 100 simulations with randomised connectivity and synaptic weights.

What processes underly the decision to swim, the start side, and the delays to the first motor response? How is synchrony avoided? To answer these questions, we study *CNS model* responses to a repeated, weak, left side trunk skin stimulus. We found that the side of first motor response depended on the interplay of three factors: 1) synaptic strengths of excitation from *exIN*s to *hdIN*s; (2) the level of *hdIN*s electrical coupling; and (3) the strength of *cIN* inhibition.

We test the hypothesis that the long and variable excitation in *hdIN*s which can lead to the start of swimming derives from the firing of *exIN*s on each side of the hindbrain (Koutsikou et al., 2018). Preliminary recordings have shown that some *exIN*s, which are normally silent at rest, fire irregularly following a 1ms trunk skin stimulation and single spike firing of sensory pathway neuron populations at short delays (<13ms; Fig. 4c). In the *CNS model, exIN*s are excited to fire by sensory pathway neurons and extend this brief sensory input by producing reverberating, irregular firing (Methods) generated via their recurrent ipsilateral and contralateral excitatory connections (Figs 4d,g-j). The *exIN*s excite *hdIN*s and the other CPG neurons, generating a ramp of glutamatergic excitation lasting up to 1.5 seconds on both body sides. This ramp can be seen in the model by averaging *hdIN* potentials in different trials in the case where the swimming is not initiated (Fig. 4f) and it qualitatively matches the average of recorded *dIN*s voltages in experiment (below swimming threshold case, Fig. 4f). If the excitation from *exIN*s is stronger on one side, the *hdIN*s are recruited to fire by this excitation (Fig. 4g) and they recruit the whole electrically coupled *dIN* population on that side (Hull et al., 2015; Li et al., 2009). This firing recruits other CPG neurons and *mn*s to produce the first motor response. As shown below, a balance in the level of *hdIN* excitation on the two sides is necessary to initiate swimming reliably.

The excitation from *exIN*s to *hdIN*s depends on their irregular firing, their connectivity to *hdIN*s and the strengths of these synaptic connections. All of these factors play a role in the decision making process. To measure the sum of their contributions, we calculate the average potentials *V*_*L*_ (*t*) and *V*_*R*_ (*t*) for left and right hdINs (Fig. 4 g-j, blue and red lines in the middle panels). If the average potentials *V*_*L*_ (*t*) and *V*_*R*_ (*t*) cross the empirically selected threshold level (−27mV) with a time difference more than 4 ms then firing of sufficient right (or left) *hdIN*s occurs to initiate swimming. The dynamics of *V*_*L*_ (*t*) and *V*_*R*_ (*t*) and the time difference between threshold crossings are indicators of the swimming starting side and time.

When stimulated on the left side there are four response types in Fig. 4 g-j:

1. **Swimming starts on the unstimulated or stimulated side (Fig. 4g):** Both *V*_*L*_ (*t*) and *V*_*R*_ (*t*) reach threshold but the time difference between threshold crossings is relatively large. For the example in Fig. 4g the potential *V*_*R*_ (*t*) grows faster, activates right CPG neurons including *cIN*s whose inhibition leads to the significant decrease of *V*_*L*_ (*t*) However, due to post inhibitory rebound of left *dIN*s, potential *V*_*L*_ (*t*) increases and crosses the threshold after a delay of about 40ms. As a result, swimming starts on the right (unstimulated) side. Similar dynamics of *V*_*L*_ (*t*) or *V*_*R*_ (*t*) with fast growing potential *V*_*L*_ (*t*) can lead to swimming starting on the left (stimulated) side (not shown).
2. **No swimming (Fig. 4h):** Neither *V*_*L*_ (*t*) nor *V*_*R*_ (*t*) reaches the threshold and swimming is not initiated.
3. **One sided activity (Fig. 4i):** Either *V*_*L*_ (*t*) or *V*_*R*_ (*t*) reaches the threshold and CPG neurons on one side fire briefly but this activation is not sufficient to initiate swimming. A similar activity pattern, probably equivalent to a simple flexion behaviour, has been seen rarely in experiments.
4. **Synchrony transition to swimming (Fig. 4j):** Both *V*_*L*_ (*t*) and *V*_*R*_ (*t*) reach the threshold with a small time difference (less than 3ms). In synchrony, CPG neurons on both sides fire simultaneously for 1-15 cycles before switching to swimming (Ferrario et al., 2018; Li et al., 2014b).

One sided and synchronous activity in simulations critically depends on an imbalance in the *exIN* to *hdIN* excitation on the two sides. It was previously shown that the generation of post-inhibitory rebound spikes in *hdIN*s requires a sufficient level of glutamatergic depolarisation (Ferrario et al., 2018; Roberts et al., 2014). One sided activity occurs when the *exIN* to *hdIN* excitation on one side is high enough to generate firing, while on the opposite side excitation is too low for firing on rebound following *cIN* inhibition. Synchrony occurs when the *exIN* to *hdIN* excitation on both sides builds up at similar rates and is strong enough to initiate a simultaneous firing of *hdIN*s on both sides.

The commissural connection probability of left and right *exIN*s has a strong influence on the motor response (Fig 4k). In experiments, low level trunk skin stimuli lead to swimming in ∼ 50% of trials. Therefore, we selected the optimal value of *exIN* commissural connection probability *p*=0.33 to obtain a similar percentage and minimize the percentage of “undesired” one sided and synchrony responses.

### OPERATION OF CNS *MODEL:* SWIMMING INITIATION

The *CNS model* can reproduce responses to trunk and head skin stimulation (Buhl et al., 2012; Koutsikou et al., 2018; Roberts et al., 2014). Stimulation of four sensory *RB*s on the left side to fire once (Fig. 5a) mimics trunk skin touch (TT). This triggers single spikes in the left side population of *dla*s and *dlc*s, which project rostrally to the hindbrain to excite *exIN*s on both sides. The excitation initiates long-lasting reverberating activity in *exINs* and summation of EPSPs in hindbrain *dIN*s reaches the firing threshold on the right side. The electrical and chemical coupling between *dIN*s then recruits the whole *dIN* population as well as other CPG neurons and the decision to swim is made. After the initial spikes on the right, CPG neurons on the left receive inhibition from *cIN*s which leads to post-inhibitory rebound (PIR) spikes in left *dIN*s which activate left CPG neurons. Thus, the anti-phase, left-right oscillatory firing of swimming is sustained by PIR and glutamatergic mutual excitation of *dIN*s.

**Figure 5:**
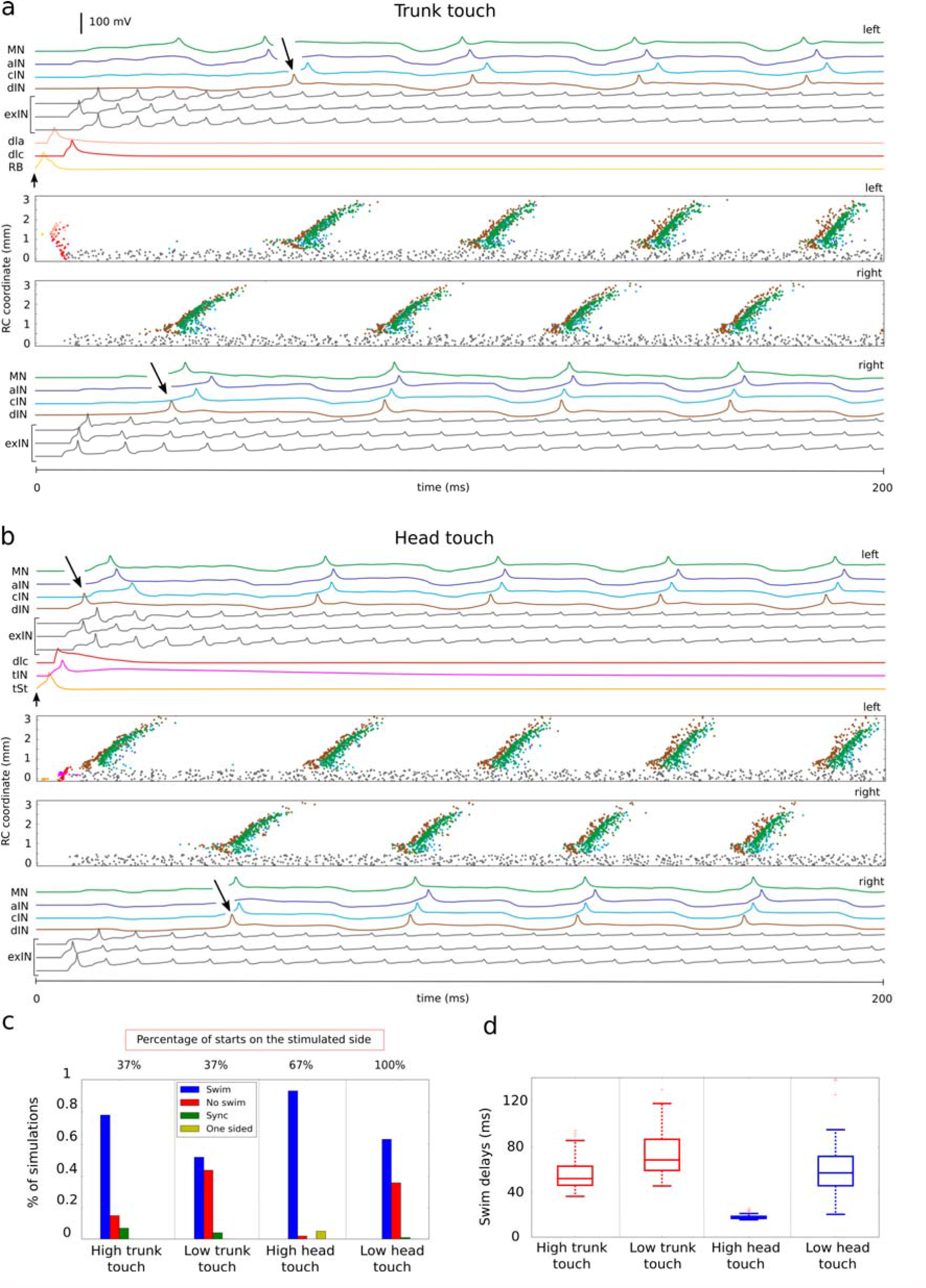
Swimming initiation in response to trunk and head touch stimulation. **a-b**. Neuronal activities on the left and right sides (2 upper and two lower panels, respectively). Central panels show spiking times where the vertical coordinate corresponds to the neuron rostro-caudal position. Other panels show the voltage traces of selected active neurons (one from sensory pathway and CPG populations plus three *exIN*s) and black arrows indicate the first *dIN* spike. **(a)** To mimic trunk touch 4 *RB*s are activated by a brief stimulation at time 0 (arrow, amplitude 0.3nA, duration 5ms). This starts swimming. **(b)** To mimic head touch, 4 sensory *tSt*s fire one spike in response to stimulation at time 0 (arrow, amplitude 0.3nA, duration 5ms). **c-d**. Summary of statistics for simulated responses to high (low) trunk or head skin stimulation. **(c)** Simulation outputs are classified as swim, no swim, synchrony (sync) and one sided. Distribution of outputs is shown for each protocol as well as the percentages of swimming starting on the stimulated side. **(d)** Boxplots of swimming delays for each type of stimulation.

Stimulation of four sensory *tSt* neurons on the left side to fire once (Fig. 5b) mimics the head skin touch (HT). They excite *tIN*s and *rdlc*s which fire once to excite *exIN*s on both sides and initiate reverberating firing. Short latency direct inputs from *tIN*s and *exIN*s recruit left dINs which then excite left CPG neurons and initiate swimming.

Simulations of *CNS model* were used to investigate the effects of the strength of stimulation and our results are in a good correspondence with experimental studies on trunk and head touch initiation with **high (low)** stimulation levels. To mimic *high* (*low*) stimulation (Buhl et al., 2015; Koutsikou et al., 2018) (also STAR Methods) we excite four (two) *RB*s for trunk and four (two) *tSt*s for head skin stimulation, respectively. Figs 5c-d shows a summary of 100 repeated simulations: **High TT** stimulation leads to a swimming start with variable delays in 78/100 cases. **Low** level **TT** stimulation generates less reliable swimming initiation with longer and more variable delays in 52/100 cases. For both stimulation levels, the percentage of swimming starts on the stimulated side was 37% (in line with experimental 32% (Koutsikou et al., 2018)). For **high HT** stimulations the swimming starts very reliably (95/100) with short delays and always on the stimulated side. **Low HT** stimulation leads to less reliable initiation (78/100) with variable delays and 67% of swimming starts are on the stimulated side.

### OPERATION OF CNS *MODEL*: SWIMMING TERMINATION

As the tadpole continues to swim, the frequency slowly drops and swimming can stop spontaneously. In life this is due to adaptive changes in the excitability of active neurons which release purinergic transmitters ATP and adenoside during swimming and reduction of inward current in *dIN*s (Dale, 2002; Li and Moult, 2012; Roberts et al., 2010a). We use a simplification to model this phenomenon, and introduce synaptic depression to recurrent *dIN* NMDA conductances. This leads to the failure of PIR spiking in the dIN population and swimming stops (Fig. 6a).

**Figure 6:**
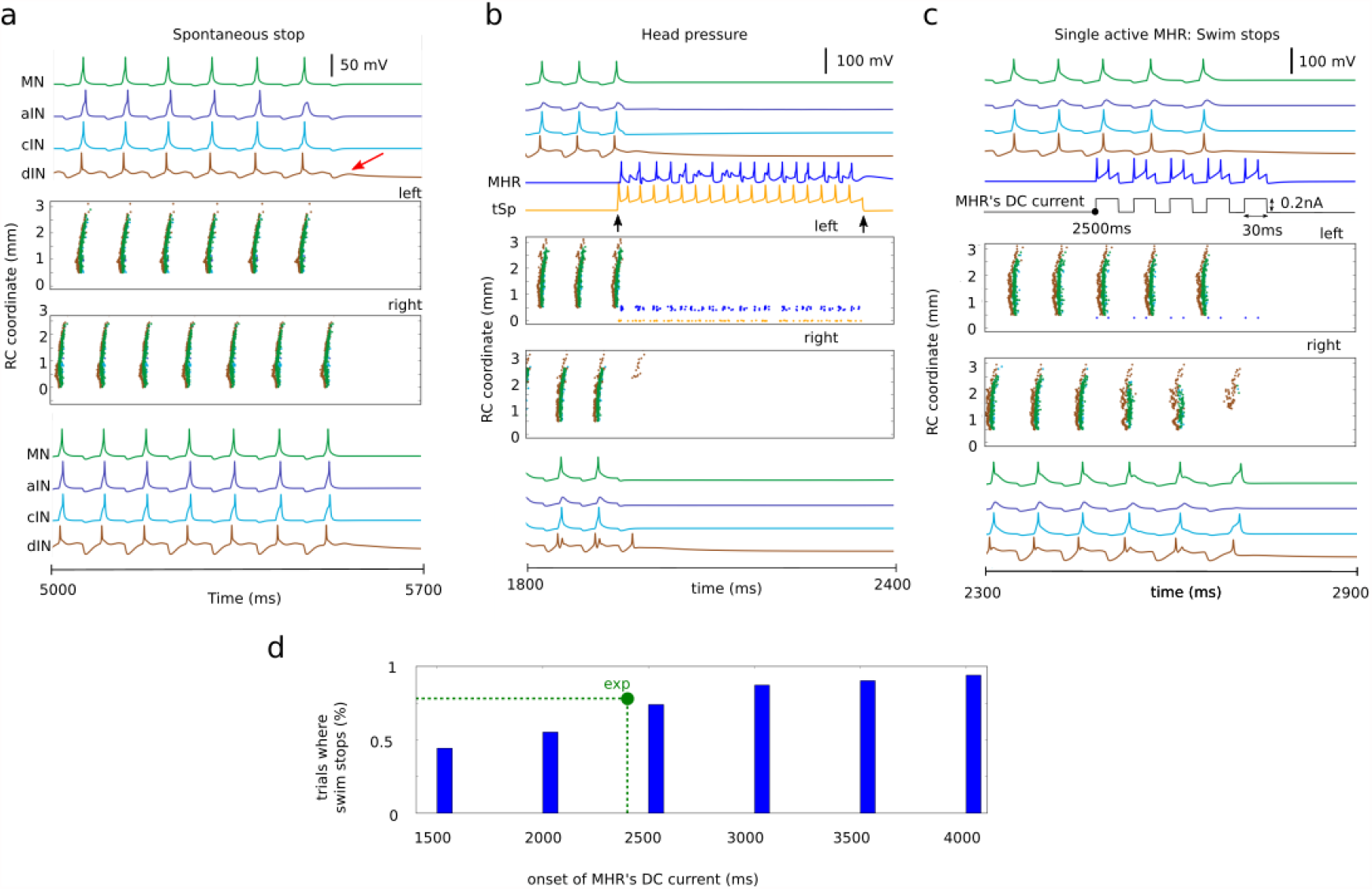
Swimming termination. **a-c**. Neuronal activities on the left (top) and right (bottom) body side for one selected simulation of the *CNS model* near swimming termination. Central subpanels show neuron spike times vs their rostro-caudal position (vertical) and upper and lower subpanels display the voltage traces for different types of CPG neurons. **(a)** Swimming slows before spontaneous termination by synaptic depression at about 5.6 s after stimulation. In the left side *dIN* trace there is no rebound spike after the last reciprocal IPSP (red arrow). **(b)** To mimic the head skin pressure, 10 sensory *tSp* neurons on the left side are injected with a step current (duration 0.4 s, 0.2nA). Both *tSp* and *MHR* neurons fire rhythmically (the mean frequency 32.5Hz), inhibit CPG neurons on both sides and stop swimming. In this case some caudal *dIN*s on the right fire after the start of the inhibition but there is no mn activity. **(c)** A single randomly selected *MHR* is injected with 5 equal DC current pulses starting at 2,500 ms (total duration 0.3 s, each pulse is 30 ms). The single *MHR* fires twice to each pulse, inhibits CPG neurons and this is sufficient to stop swimming. **(d)** Percentage of simulations where swimming was stopped by the activation of a single randomly selected *MHR* vs the onset time of the *MHR*’s current, green dot shows experimentally determined value of stoppings for 2.4 s onset.

The tadpole usually stops when its head bumps into something and it attaches with mucus from the cement gland (Perrins et al., 2002) (Fig. 1a). To model swimming termination by head skin pressure we inject current to evoke multiple firing in 10 head pressure sensory *tSp* neurons on one side (Fig 1c). Their firing activates all ipsilateral *MHR*s which project axons to both sides and generate long-lasting GABAergic inhibition of CPG neurons to terminate swimming (Fig. 6b). Repeated simulations with different connectivity and synaptic strengths show reliable termination in all cases.

Experiments have shown that the stimulation of single *MHR* after 2.4 seconds of swimming can stop swimming in 78% of trials (Perrins et al., 2002). To mimic this experiment, we stimulate one randomly selected *MHR* neuron 2.5 seconds after the initiation of swimming in the CNS model (Fig. 6c). Swimming stopped in 74% of 100 trials with different connectomes. Further simulations showed that stopping reliability increased monotonically with increasing time during a swimming episode due to decreasing reliability of *dIN* spike generation (Fig. 6d), green dot shows experimentally determined value and allows to compare experimental and modelling findings.

## BIOMECHANICAL *VT MODEL*

Biomechanical modelling of tadpole swimming is based on the “Sibernetic” software system (Palyanov et al., 2016) which has been used successfully to model the 3D body of the “worm” *C. elegans* and simulate its swimming and crawling locomotion (Palyanov et al., 2018; Sarma et al., 2018). In Sibernetic, objects and their environment are composed of equally spaced and sized “*particles*” interacting with each other. They can be of different mass to represent liquids or other materials with different densities. Elastic objects are constructed by connecting “*particles”* with “*springs*” with individually defined stiffness and, for muscle tissue, groups of user-defined springs can be instructed to contract in response to incoming motor signals from the *CNS model*. Fluid dynamics calculations are based on the predictive-corrective incompressible smoothed particle hydrodynamics algorithm (Solenthaler and Pajarola, 2009) which considers viscosity, surface tension, density and pressure, as well as gravity and other external forces acting on a liquid.

Reconstruction of the 3D body of the tadpole used photographs, scale drawings and histological sections (Fig. 7 a-c). It reproduces its surface shape, length, width, and height and also internal structures: notochord (a longitudinal rod providing stiffness (Adams et al., 1990)), segmented swimming muscles, undifferentiated belly and cement gland on the head which is adhesive to solid surfaces (Fig. 7a-d). Striving for a balance between model accuracy and computational performance the model body has 14,005 *particles* and is 100 *particles* in length. By adjusting the properties of *“springs”* connecting *“particles”* it is possible to define the stiffness coefficients of various parts of the body. For example, the notochord is more rigid, while the belly region is less rigid and denser.

**Figure 7:**
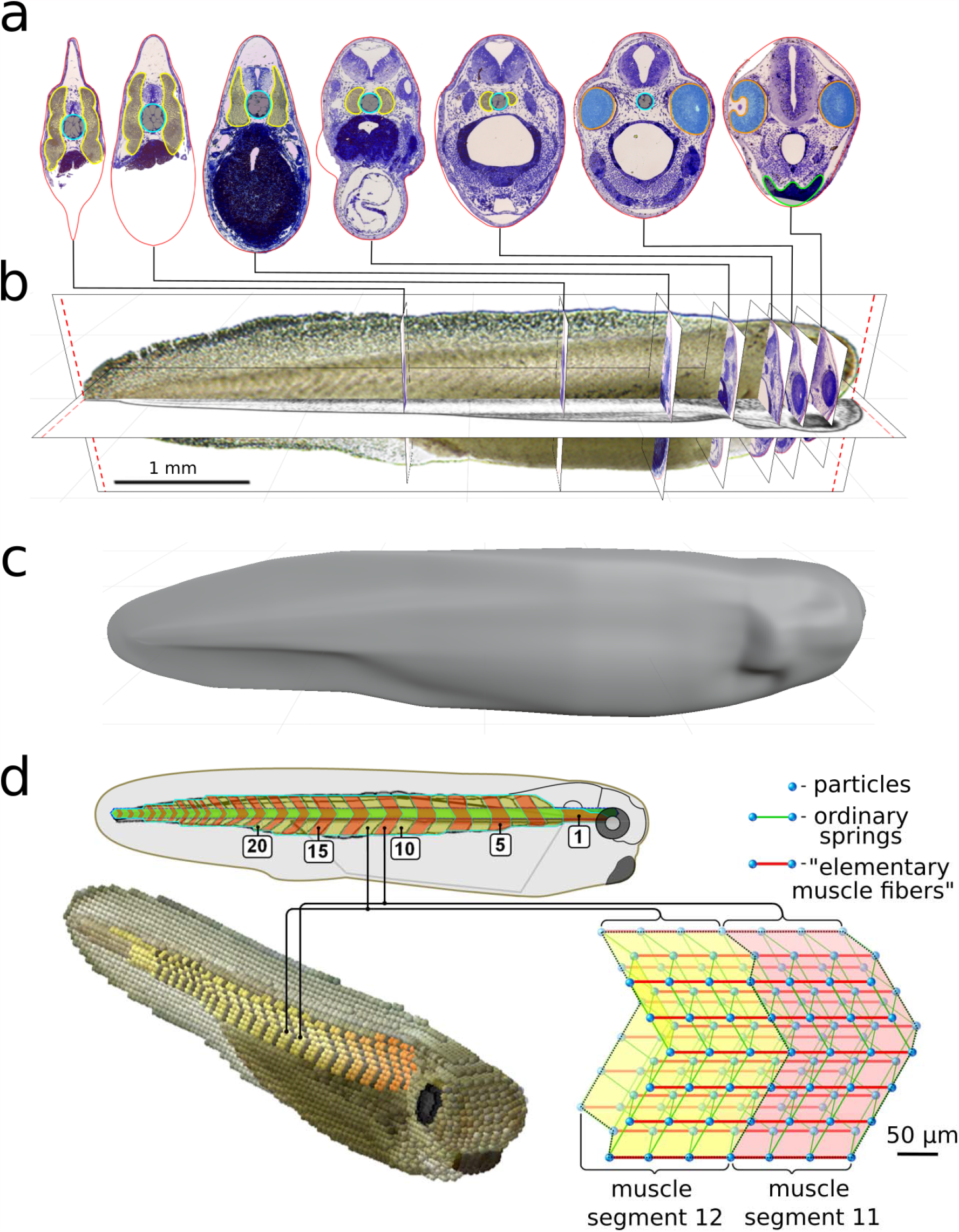
*VT model:* 3D body reconstruction and swimming in “water”. **a-c**. Reconstruction of 3D tadpole body from experimental images and body sections of the real tadpole (using the Blender software). Some body sections at different RC positions from head to tail are shown in **(a)**. **d**. Scale diagram of a real tadpole (top) with the notochord (green) and muscle segments (yellow and orange) with a schematic representation of muscle segments (11 and 12) in Sibernetic (bottom-right). This simplified 3D construction includes particles (blue balls) connected by ordinary springs (green) and “elementary muscle fibers” (red) which can contract in response to motoneuron spiking.

We took care to reproduce the geometry of the segmented muscles that control the body and generate swimming (Fig. 7d). Muscles contain longitudinal contractile “*springs”* (elementary muscle fibers: ELMs) with contraction dynamics based on adult frog muscle. *Particles* all over the muscle surface connect with *particles* of adjacent tissues including the notochord. Segments can contract independently when activated by signals from *CNS model* motoneurons (Fig. 7d, 8a). To observe swimming the *VT model* of the tadpole body is placed into a tank filled with “water” (Fig. 8).

**Figure 8:**
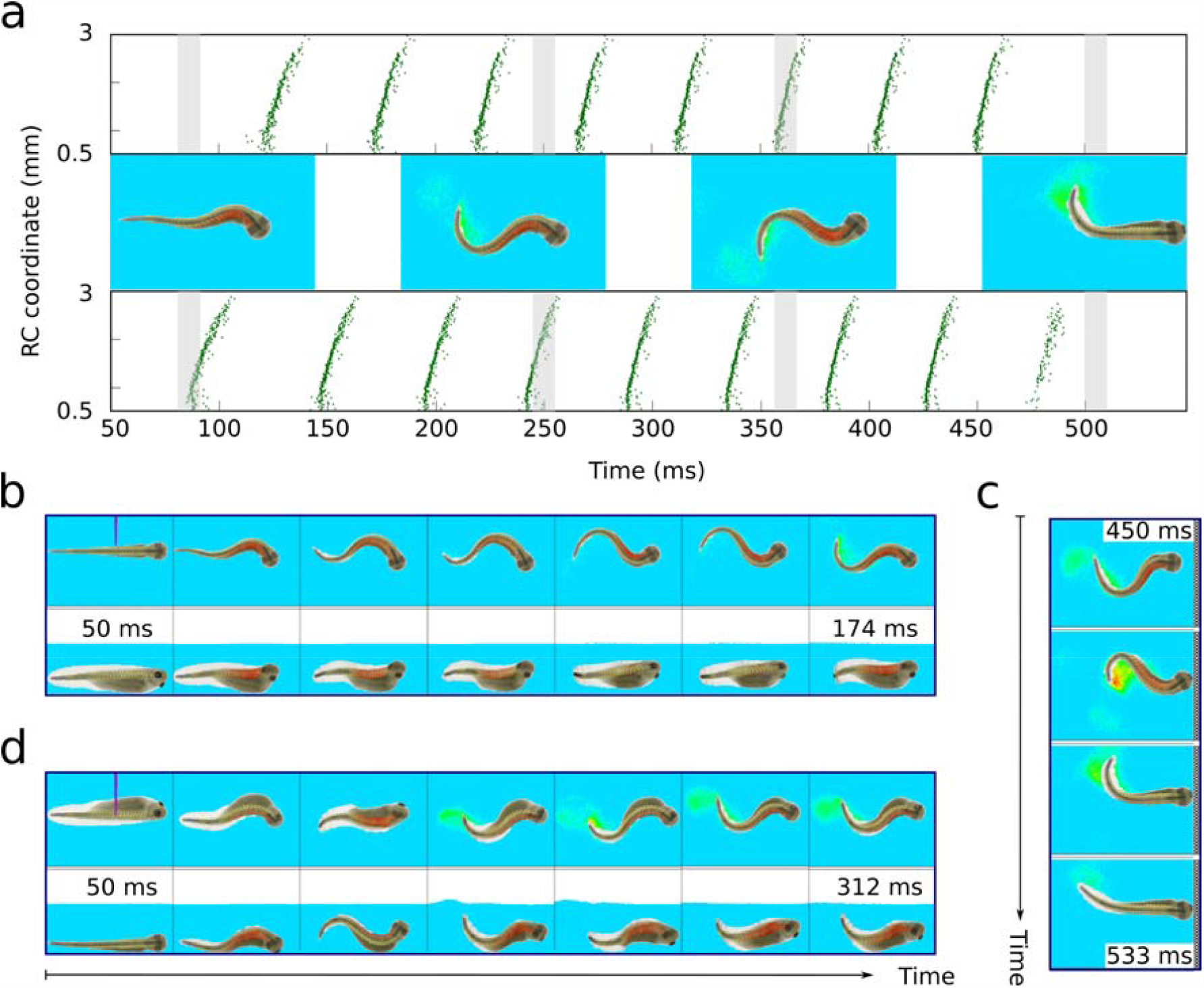
Motoneuron spiking, muscle contraction, swimming and stopping. **a**. *CNS model* motoneuron spiking and corresponding body positions during swimming. Green dots show motoneuron spiking times. Horizontal and vertical axes show motoneuron spike time and RC position. Spike related body shapes are in the middle and grey strips show correspondence between spiking and *VT model* swimming (each body shape relates to the middle time point of grey strip). The first frame shows active rostral muscles (red) for right flexion initiation. Next two frames show right and left flexions and the last frame corresponds to stopping. Body-water interaction is visualised by speed of water particles (blue-yellow-red corresponds to slow-medium-fast movement). **b**. Stimulation on the left side at 50 ms follows by flexion on the right and left sides. Frames are from video1, upper and lower panels show top and right side views, respectively (in **b** and **d**). **c**. Sequence of frames from video1 shows the body movements near the swimming stopping. The top frame shows the tadpole head approaching the wall and the next frame corresponds to the wall touch and mucus release. Last two frames show the attached head and tail moving by inertia. **d**. Frames from video2 show the body rotation during swimming. At initial (left) frame the tadpole lies at the bottom on right body side and left side is stimulated. Next frame shows the first right side flexion and movement start. Eventually, the body orientation changes and two of the most right frames show tadpole swimming in dorsal-up position in the middle of tank.

We project a spatio-temporal pattern of motoneuron spiking activity generated by *CNS model* during swimming to the muscle segments on each side of the body (Fig. 8a). Each CNS model motoneuron spike at time *t*_*spike*_ causes each ELM within an innervated muscle segment to contract with a force described by the “bell-shape-type” function centred at time (*t*_*spike*_ + 20) ms with summation of forces generated by nearest spikes (Fig. 8a and Methods), where 20ms is the delay between motoneuron spike and the maximal ELM force. One side flexes first (contracting muscles are red) and the body interacts with the “water” where the velocities of liquid particles are visualized. Alternating flexions then follow and the VT tadpole generates realistic swimming (Supplementary video1). Key frames from video1 show body movements and the corresponding spiking times of motoneurons distributed along the body (Fig. 8a). These illustrate swimming initiation and stopping. Initially, the tadpole rests in the middle of the tank near the bottom, the left trunk is stimulated and swimming starts with flexion to the right side (Fig 8b). Swimming stops when the head touches the wall and *VT model* sticks with cement gland mucus (Fig. 8c).

In video2 the tadpole initially lies on its right side on the tank bottom. Skin stimulation on left leads to muscle contraction on the right side (Fig. 8d) and, as swimming continues, the body rotates to a standard dorsal-up, belly-down position. As in real tadpoles the *VT model* moves slowly upwards in the “water” as it swims and, if the tadpole lies on its left or right side and then starts swimming, it is able to reach a dorsal-up position in 2-4 cycle due to a simple belly-ballast effect (Roberts et al., 2000).

Bottom-left: Reconstructed 3D tadpole body (*VT model*) in the Sibernetic system. The 3D shape of the tadpole body **(c)** was loaded and visualized by Sibernetic to assist in building of the *VT model*. The density of the *VT model* belly is higher (1060 kg/m^3^) than other tissues (1035 kg/m^3^). To simulate swimming the *VT model* is placed into a tank (18.4×3.7×3.7 mm) filled with water represented by ≈ 2 billion liquid particles with density and dynamical viscosity (measured in simulation using Stokes’ law) equal to those of water at 20°C. Swimming speed of the *VT model* ≈ 19.3 mm/s (tadpole length is 5 mm and period of muscle contraction on one body side is 100 ms), whereas the typical preferred speed of a real tadpole is ≈ 21 mm/s. The difference might be due to the ciliary activity leading to tadpole gliding which is not included to the model (Hoff and Wassersug, 2000).

## DISCUSSION

The *CNS model* is a detailed and biologically-realistic reconstruction of the tadpole nervous system and is formed by a spiking neuronal network that reproduces the dynamics of neuronal responses seen in electrophysiological recordings. The model receives and integrates sensory signals from three skin sensory pathways to make decision and control swimming initiation and stopping.

The spiking activity of motoneurons generated by the *CNS model* is used to activate muscles in the biomechanical *VT model* which then generates body movements very similar to those of the real tadpoles swimming in water. Our results present a first attempt to build a detailed 3-dimensional model of a whole animal’s body and show how its locomotor behaviour is controlled by CNS neuronal networks.

### Building model circuits

The *CNS model* includes neuronal circuits related to different functions. Finding pair-wise connectivity inside and between circuits is a difficult problem and our results build on a growing literature on recent developments in the network neuroscience (Bassett et al., 2018). Our methods combine techniques from probability theory with anatomical data to constrain and define connections. Our anatomical modelling considers the independent Bernoulli variables and similarities in connections between different neuron populations. This method highlights a path to a general technique for finding pairwise connectivity in complex circuits.

### Variable spiking activity but reliable function

The general view on spiking level modelling in neuroscience states that spike trains generated by neuronal circuits are highly variable while the function of the circuit is stable and reliable (Abbott et al., 2016; Deco et al., 2015). The *CNS model* includes several sources of variability and the CNS model generates variable spike trains. Remarkably, mapping these spike trains to the *VT model* shows reliable results in swimming behaviour similar to real tadpole movements.

### Decision making, variability and unpredictability

Several theoretical ideas on decision making are under discussion (Schall, 2001; Wang, 2012). However, it becomes increasingly clear that detailed computational modelling (Lo and Wang, 2016; Shevtsova et al., 2015) which allows comparison with experimental recordings is rather limited. Our *CNS model* with realistic connectivity and neuron- and synapse-level resolution explains how the decision to swim is made: irregular reverberating firing of sensory memory *exIN*s produce the long and variable ramps of excitation in reticulospinal hindbrain *dIN*s on both sides which are coordinated by commissural *exIN* connections and determine the side of first flexion and the initiation of swimming. It is known that there are many similarities in the organization of neuronal circuits controlling locomotion in vertebrates (Kiehn, 2016) and our simulations shed a light on the basic neuronal processes of decision making for movement initiation in all vertebrates.

### A new approach to modeling

The study of motor control and behaviour is usually centred on neuronal networks for signal processing. It is known that motor behaviour depends on interactions between the nervous, musculoskeletal systems and the external environment (Beer, 2009; Tytell et al., 2011; Williams and McMillen, 2015). Our “whole animal” combination of detailed neuronal and biomechanical models explains neuronal mechanisms and reproduces realistic swimming in water.

## STAR METHODS

### Anatomical modeling

Each neuron in the CNS model is characterized by a type and a RC coordinate (measured in microns from the midbrain-hindbrain border). We have previously shown how to define the locations of tadpole neurons and their connectivity using a “developmental” method (Borisyuk et al., 2014) that simulated the growth of axons and synaptic formation based on anatomical measurements (Roberts et al., 2014). We use this method for CPG [*dIN, cIN, aIN, mn*] and sensory pathway populations [*RB, dla, dlc*].

For the remaining neurons [*tSt, tSp, tIN, MHR, exIN*] we use anatomical measurements and uniformly distribute RC coordinates of sensory pathway neurons in the following intervals: *tSt* in [−50,0] *μm*(Buhl et al., 2012); *tSp* in [0,10] *μm* (Hayes and Roberts, 1983); *tIN* in [150,330] *μm* (Buhl et al., 2012); *MHR* in [400,550] *μm* (Perrins et al., 2002). RC coordinates of hindbrain sensory memory neurons *exINs* have been uniformly distributed in the caudal part of the hindbrain in range [− 130,500] *μm* (hypothetical assumption based on indirect data (Koutsikou et al., 2018).

To find connections to and from these cell types we use a probabilistic approach (Ferrario et al., 2018), where the directed connection from neuron *i* to neuron *j* is represented by the Bernoulli variable *X*_*ij*_ with probability *p*_*ij*_= *pr*{*X*_*ij*_ = 1}.We constructed connection probabilities by generating 1000 connectomes using the developmental approach and averaging across connectomes. We use these probabilities to infer the connectivity between populations where there is limited experimental data by hypothesizing anatomical and functional similarities. For example, connections probabilities from trunk skin sensory *RB*s to sensory pathway *dla*s are high, similar to connections from head skin sensory *tSt*s to sensory pathway *tINs*. This similarity has been shown by direct pairwise recordings (Buhl et al., 2012). Using this similarity, we extrapolate from the known connection probabilities of *RB*s to *dla*s in our developmental model (Roberts et al., 2014) and define connection probabilities for *tSt*s to *tINs* connections (Algorithm 1).

In many cases some anatomical characteristics like the RC positions of neurons are known. We use this information and define the conditional probability of connection given the distance between neurons (Algorithms 2). For example, for *tINs* to *dINs* connections are inferred based on the similarity with *RBs* to *dlas* connections and additionally by considering the RC distances between *tINs* and *dINs*.

### Algorithm 1: Anatomical similarity and generalization procedure

Assume that there is a pair of populations S1 with known probabilities of pairwise connections from the first to second population. Consider another pair of populations S2 with unknown connection probabilities. Also, we assume that experimental evidence allows us to propose a similarity between connectivity in pair S1 and in pair S2. After that we use a generalization procedure (Borisyuk et al., 2014): we estimate the cumulative distribution of known connection probabilities in S1 based on the developmental model and use this distribution to randomly prescribe a probability to each connection in S2.

### Connections *tSts->tINs*

The results (Buhl et al., 2012) provide direct evidence of high probabilities of connections from *tSt*s to *tIN*s. We expect that these connections and connections from *RB*s to *dla*s are similar because in both cases we consider the ipsilateral projections from sensory neurons (*tSts* and *RBs*) to sensory interneurons (*dla*s and *tINs*) and probabilities of connection from *RB*s to *dla*s are relatively high. Thus, for known probabilities of connections from *RBs* to *dlas* we calculate the piecewise approximation of the cumulative distribution of probabilities and use it to randomly prescribe connection probabilities from *tSt* to *tIN* neurons.

### Connections *tSts->rdlcs*

Similarly, results (Buhl et al., 2015) provide data for similarities of *tSt* to *rdlc* connections and *RB* to *rdlc* connections: both connections are reliable, ipsilateral and from sensory neurons (*tSt*s and *RB*s) to *dlc*s. Additionally, in both cases the connections have similar functional properties (provide the sensory signal to initiate swimming). The generalization procedure is then used to find individual connection probabilities.

### Connections *tSps->MHRs*

Previous results (Perrins et al., 2002) provide indirect evidence that these connections have a relatively high probability like connections from *RB*s to *dla*s, and in both cases connections are ipsilateral from sensory cells (*RBs* and *tSps*). Thus, we assume similarity and use the generalization procedure to extract the *tSps* to *MHRs* connection probabilities from *RB* to *dla* probabilities.

### Algorithm 2. Similarity + additional anatomical data

Algorithm 2 uses similarity between connections in two pairs of populations (like in Algorithm1) and additionally the probability distribution of axonal lengths to find connection probabilities. <u>Assume</u> that S1 is a pair of populations with known probabilities of connections and consider another pair of populations S2 with unknown connection probabilities. Also, we assume that experimental evidence allows us to propose a similarity between connectivity in pair S1 and in pair S2. <u>Additionally</u>, we consider the known probability distribution of presynaptic axons in S1 and assume that the presynaptic axonal lengths in S2 have a similar probabilistic distribution.

The algorithm:

1. Assume that all neurons in each pair (S1/S2) are ordered according their increasing RC coordinate.
2. For each ordered pair of neurons (*i,j*) in S1 consider the connection probability *P*_1_ and the Euclidean distance *L* between RC coordinates of these two neurons. We then extract the dependence of the probability of their connection on the distance between the two neurons: *P*_1_ = *f*(*L*)
3. For each ordered pair of neurons (*m,k*) in pair S2 find the distance *Q* between RC coordinates and use the constructed function *P* = *f*(*L*)to find the probability of connection from neuron *m* to *k*: *P*_2_ = *f*(*Q*).

### Connections *tINs->hdINs*

We know from experiments that *tIN*s have a relatively long descending axons: the mean is 1750 *μm* (s.d. is 480) (Buhl et al., 2012) and similar, for ascending *dla*s axons the mean is 1820 *μm* (s.d. is 470) (Li et al., 2001). Additionally, both *tIN* and *dla* neurons have dorsally located dendrites and fire transiently in response to a touch (either head or trunk skin) and excite CPG neurons (Buhl et al., 2012; Li et al., 2007; Li et al., 2004b). Thus we expect *tIN*s and *dla*s to have similar connection probabilities with CPG neurons. We use the *dla* to *cIN* connection probabilities to define the function *P* = *f*(*L*),where *L* is the RC distance between coordinates of *dla*s and *dIN*s and we are taking into account that *dla*s are ascending but *tIN*s are descending neurons; therefore we consider the reflection symmetry for calculation distances. Using this constructed function we find the connection probabilities for *tIN*s -> *hdIN*s.

### Inhibitory connections from *MHR* to CPG neurons

(Perrins et al., 2002) provides indirect evidence that reticulospinal *MHR* neurons connect with high probability to CPG neurons in the rostral part of the spinal cord, sending a powerful inhibitory signal that stops swimming. We use similarities between connections from *dlc*s to all CPG populations and connections from *MHR*s to all CPG populations. There are two reasons for doing that: First, the axons of *MHR* neurons are relatively long, the mean length is 1180 *μm* (s.d. is 350) (Li et al., 2001) and the length distribution is similar to the distribution of *dlc* axon length (the mean length is 1190 *μm* (s.d. is 410) (Li et al., 2001). Second, both *MHR* and *dlc* neurons have a similar functional role in the swimming network, they fire transiently in response to sensory input and connect to CPG neurons. We therefore expect that both *dlc*s and *MHR*s to have similar connection probabilities to CPG neuronal populations.

We apply Algorithm 2 to find connections from *MHR*s to each of four CPG populations (*dIN, cIN, aIN* and *mn*). We construct four functions *P* = *f*(*L*) and use them to establish probabilities of contralateral connection from *MHR* to each of four CPG subpopulations. As *dlc*s are ascending and *MHR*s are descending neurons, we apply the reflection symmetry to function *P* = *f*(*L*). We know that *MHR* neurons have primary contralateral axons, and about 20% of *MHR*s have secondary ipsilateral axons (Perrins et al., 2002). Thus, we randomly select 20% *MHR*s in the model and use the same functions *P* = *f*(*L*) to define probabilities of ipsilateral connections from *MHR*s to the corresponding CPG neuronal population.

### Probabilities of connections from *dla* (*dlc*) to *exIN*

Data on *exIN* anatomy is not yet available and electrophyisiological data is limited (Roberts et al., 2019). Preliminary experiments (Koutsikou and Soffe, unpublished) recorded the intracellular activity of 33 midbrain interneurons (*mIN*s) in response to trunk skin stimuli near threshold which initiated swimming in 50% of trials. These recordings showed that 45% of *mIN*s (15/33) showed short latency EPSP less than 13ms from stimulation. We assume that: (1) *mIN*s are analogous to the hindbrain *exIN*s because both neuronal populations generate irregular spiking activity in response to stimulation; (2) 45% of *exIN*s demonstrate early EPSPs in the interval [0, 13] ms. It is known that sensory projection neurons *dla* and *dlc* fire only once in response to sensory *RB* cell stimulation and only these neurons spike in 13 ms after the skin stimulation (Koutsikou et al., 2018; Li et al., 2003; Li et al., 2004a). Therefore, we conclude that the early *exIN* EPSPs result from synapses made by *dla*s/*dlc*s (Roberts et al., 2019).

We assume that the connection probability from a *dla* (*dlc*) neuron to *exIN* is the independent Bernoulli variable with the probability *p*_1_ (*p*_2_), respectively. The EPSP in *exIN* can appear as a result of at least one spike incoming from *dla*s or at least one spike incoming from *dlc*s. In addition, from simulations of the spinal cord model (Roberts et al., 2014) we find that the average number of these neurons active after skin trunk stimulation (stimulus to two *RB*s) is *n*_1_ = 10.4 *dla*s and *n*_2_ =22.9 *dlc*s.

Also, we assume that the probabilities of direct influence from *dla*s and *dlcs* to *exINs* are independent and equal:

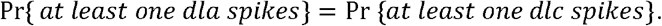

Let us consider the event A: {Appearance of EPSP in *exIN* neuron during the time interval [0, 13] ms after the stimulation}.

From the experimental data described above we know:Pr{A}=0.45.

The probability of spikes incoming to *exIN* from *n*_1_ active *dla*s can be described by the Binominal distribution *B*(*n*_1_,*p*_1_) (*B*(*n*_2_,*p*_2_), respectively, for spikes from *dlc*s).

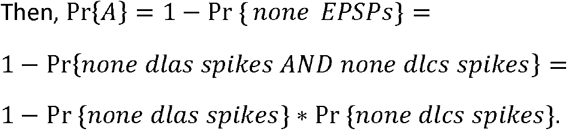

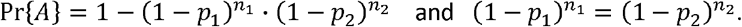

From these two formulas:

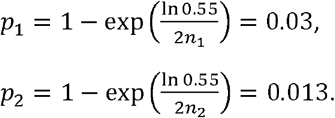

Thus, we select connection probability 0.03 for *dla*s->*exIN*s connections and 0.013 for *dlc*s->*exIN*s connections.

### Connections *tINs->exINs*

We hypothesize that *tIN*s make connections to an undefined population of hindbrain neurons *exIN*s (Koutsikou et al., 2018). Sensory pathway *tIN*s lie in the rostral hindbrain and have descending ipsilateral axons extending through hindbrain neuron into the spinal cord (Buhl et al., 2012). Both *dla*s and *tIN*s are excitatory sensory pathway neurons with ipsilateral axons and stimulation of an individual *dla* can initiate swimming (Li et al., 2004b). Thus, we assume similarity between the *tIN*s->*exIN*s and *dla*s->*exIN*s connectivity and prescribe connection probability 0.03 for *tIN*s->exINs connections.

### Establishing the *exIN* recurrent connections

We have proposed that *exIN*s are able to generate sustained reverberating firing in response to sensory input as a result of excitatory recurrent connections within the *exIN* population (Koutsikou et al., 2018). We assume that *exIN*s are located in a relatively compact longitudinal region of the hindbrain with make local connections. The population of motor neurons (*mn*) has mainly short axons and mostly local connectivity within their populations (Ferrario et al., 2018). Therefore, from the connection probability matrix of internal *mn*s connections we select 30×30 sub-matrix (30 first rows and columns because the most-rostral *mn*s are close to the hindbrain region where *exIN*s are presumably located) and use this sub-matrix to prescribe the connection probabilities for inter-neuronal connections of the *exIN* population of 30 neurons. To allow sustained activity within *exIN* population in response to input from *dla*s/*dlc*s it was necessary to multiply these probabilities by a factor of 2.5.

Preliminary simulations of the CNS model showed unbalanced excitation of CPG neurons on either side, which would not allow a coordinated initiation of swimming. In addition to ipsilateral connections we therefore added contralateral connections between the left and right *exIN* populations. In addition to the probabilities defined by the 30×30 probability matrix described above, for each pair of *exIN*s on the opposite sides we define the commissural connection with the probability *p*. We found that the value of this probability contributed to sustaining firing (with similar frequencies) on both sides and to a coordinated initiation of swimming. We therefore explored the space of commissural connection probabilities (see Results) and selected an optimal value *p*=0.33.

### Connections from *exINs* to hindbrain CPG neurons

We assume that the probability of connection from *exIN* to any CPG neuron located in the hindbrain is *p* = 0.05. This probability was adjusted to match the experimentally recorded wave of excitation in *hdIN*s when swimming was not initiated (see Results and Fig 4).

### Conductance based modelling of neuronal activity

The *CNS model* includes conductance-based single-compartment spike generation neuron models of Hodgkin-Huxley type with synaptic and axonal propagation delays and an additional 2-compartment *exIN* hindbrain neuron class (Koutsikou et al., 2018). Synaptic connections are established according the generated adjacency matrix and in addition to the chemical synapses, we also include electrical coupling between *dIN* axons. To define parameter values of electrical coupling we follow experimental data (Li et al., 2009; Roberts et al., 2014) and modelling (Hull et al., 2015) that suggested that these electrical connections are an important functional property of the *dIN* network. Most parameters of the physiological model are based on available experimental data. For example, synaptic strengths, membrane channel conductance and neuron capacitances of CPG neurons are mostly based on experimental results and then randomised according to a Gaussian distribution. All simulations were performed using NEURON 7.3 (Carnevale and Hines, 2006) with a fixed time-step of 0.01 ms.

To simulate the activity in the generated connectomes we represent each cell as a single compartment conductance based neuron of Hodgkin-Huxley type. All equations and parameter values for CPG neurons are the same as in paper (Ferrario et al., 2018). Here we provide some key model features.

The equation governing the membrane potential *V*_*i*_ for each neuron *i* (except *exIN*s) is

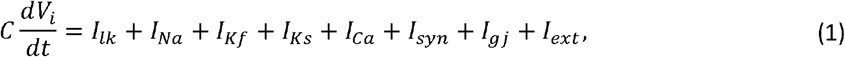

where *C* = 10*pF* is the capacitance of each neuron and corresponds to a density of 1.0 *μF* /*cm*^2^ for a total surface area of 10^−5^ *cm*^2^. *I*_*syn*_ and *I*_*gj*_ represent the sum of synaptic and gap junction inputs, respectively. The gap junction input is included only in the *dINs*, based on our previous model (Hull et al., 2015; Roberts et al., 2014). The *dIN* gap junction coupling is local and bidirectional by connecting pairs of *dINs* with rostro-caudal positions within 100 *μm* from each other. For *dIN* with index *i* the gap junction current is calculated using a simple ohmic relationship:

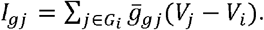

Here *G*_*i*_ is the set of indexes of all *dINs* on the same side of the body as neuron *i* located within 100 *μm* from its rostro-caudal position. The gap junction conductance 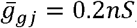, which fits experimental data on *dINs* (Roberts et al., 2014).

The term *I*_*ext*_ represents an externally injected current. All cell types except *dIN*s have no calcium component (*I*_*ca*_ = 0). The leak (*lk*), sodium (*Na*), slow and fast potassium (*Ks* and *Kf*) ion currents are given by equations (2-4):

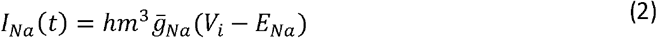

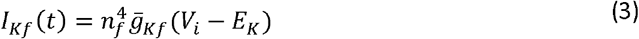

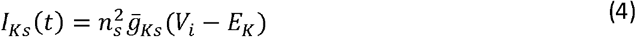

Parameters *E*_*lk*_,*E*_*Na*_ and *E*_*K*_ correspond the reversal potential for the leak, sodium and potassium channels respectively, and the parameters 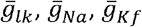 and 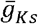 correspond to their maximum conductance. The gating variables, *h,m,n*_*f*_,*n*_*s*_ follow equations (5-8), where *X* = *h,m,n*_*f*_,*n*_*s*_.

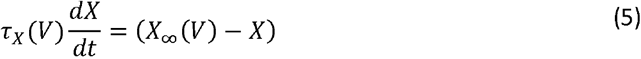

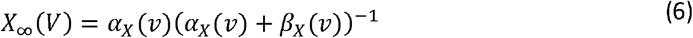

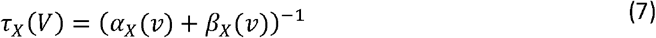

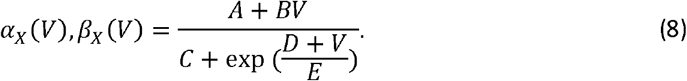

A fixed set of parameters 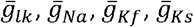 (Supplementary Table 1) and *A, B, C, D, E* (Supplementary Table 2) define the model of each cell type. Some of these parameters were reported in previous works but we included them here for completeness. Here is a brief description of the parameters and model used for each cell type.

### *RB, dla, dlc, cIN, aIN, mn* neurons

Basing on our previous developments (Ferrario et al., 2018; Roberts et al., 2014) we selected the same model and parameters for all these neuronal types. This model captures the basic properties shared by all these cell types: firing repetitively to depolarizing current.

### *dIN* neurons

The model *dIN* is based on a recent multi-compartment model (Hull et al., 2015; Hull et al., 2016). For computational limitations we simplified this model by considering only one soma/dendrite compartment. The *dINs* contain a calcium-mediated ion current in the voltage equation (1) modeled according to the Goldman-Hodgkin-Katz formulation:

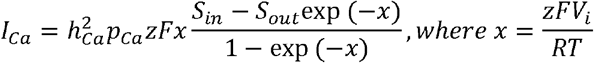

Parameter *p*_*ca*_ is the permeability of the membrane to calcium ions and parameter *z* is the calcium ionic valence (+2).*S*_*in*_ and *S*_*out*_ correspond to the calcium concentrations in and outside of the cell, respectively. Parameters *F, R* and *T* are Faraday’s constant, the ideal gas constant and the temperature, respectively. Parameters of the calcium current are: *p*_*ca*_ =14.25 *cm*^3^ /*ms,F* = 96485 *C*/*mol,R* =8.314*J*/(*K mol*),*T* = 300 *K*,[*Ca*^2 +^]_*I*_ = 10 ^−7^ *mol*/ *cm*^3^,[*Ca*^2 +^]_*o*_ = 10 ^−5^ *mol*/*cm*^3^. Finally, *h*_*Ca*_ is the gating variable associated with the calcium current, which is governed by the gating equations (5-8).

### *RB/tSt/tSp* neurons

The *RB* model is based on previous work and was modified to match the basic electrophysiological properties of these cells (Sautois et al., 2007). Detailed electrophysiological data characterizing both *tSt* and *tSp* neurons are not known. Since *RB, tSt* and *tSp* are all sensory cells we used the same *RB* model for all of them.

### *tIN* neurons

To model *tIN* neurons, we match some of the basic electrophysiological properties of these neuronal type from data in paper (Buhl et al., 2012). We start from the parameter values of the *aIN* model (Sautois et al., 2007) because both *aINs* and *tINs* have similar firing properties: repetitive firing to current injection at any depolarization level, no firing adaptation, no delayed firing and no after-spike depolarization block. We then adopt the following parameter changes:

- Leak conductance and leak reversal potential are set to match the average input resistance *R*_*in*_ = 459*M Ω* and resting potential V_rest_ = −55mV recorded in experiments (Buhl et al., 2012).
- We use the approach discussed in paper (Roberts and Tunstall, 1990) to obtain a qualitatively similar current threshold for firing. Specifically, we decrease by 25mV parameter D defining the rate functions in all ionic channels to obtain current threshold of ∼50 p*A* similar to *tINs* (from Figure 6D (Buhl et al., 2012)).
- To obtain a similar dependence of firing frequency in response to current injection (I-f curve) and match the range of firing frequencies of *tINs* we modified maximal conductance of the sodium, fast potassium and slow potassium.
- Parameter A of the *β* _*m*_ sodium rate function is decreased by 2*ms* ^−1^ to give a better match of the voltage amplitude of spikes during high current injections.

### *MHR* neurons

We model *MHR* neurons to match the electrophysiological properties of these cells (Perrins et al., 2002). We start from *dlc* model parameters (Sautois et al., 2007) because the *dlcs*’ resting potential and input resistance are closer to the average *MHR* values than all other neuronal types in the swimming circuit. Moreover, *MHR*s have similar firing properties to *dlcs*: multiple action potential firing in response to positive current injection and firing adaptation (Perrins et al., 2002; Roberts and Sillar, 1990). We then adopt the following changes:

- The leak conductance and leak reversal potential are set to match the average input resistance *R*_*in*_ = 262*M Ω* and resting potential V_rest_ = −68mV of experimentally recorded *MHRs*. These values were measured from sharp microelectrodes, which record more negative resting potential values than more precise patch electrodes. To correct these imprecise measurements, we select a lower value of the resting potential: *V*_*rest*_ = −60mV. This change was suggested by multiple comparisons between neuronal recordings made using both types of electrodes (microelectrodes and patch clamp).
- We decrease parameter D defining the rate functions of all ionic channels by 10mV to obtain a similar current threshold for firing to *MHRs*, following the same approach discussed for *tINs*.
- We match the I-f curve and firing frequencies of *MHRs*. To obtain multiple firing at any level of injected current, lower the spike frequency to physiological values and avoid single-spikes to current injection parameter A of the *α*_*m*_ sodium rate function was decreased by 3*ms* ^−1^ and A of the *β* _*f*_ of the fast potassium rate function was decreased by 0.9*ms* ^−1^.

### *exIN* neurons

The model *exIN* is based on the one proposed in paper (Koutsikou et al., 2018). Since few experimental recording of these cells are available, parameters values are randomly chosen from a physiological range to obtain repetitive firing patterns similar to the ones observed in responses to trunk and head touch. The *exIN* model is has two compartments: a combined dendrite and soma compartment and an axon compartment. The equations governing the dynamics of both compartments are based on the same Hodgkin-Huxley equations (equation (1)) with parameter values of *mn* cells (Koutsikou et al., 2018; Roberts et al., 2014; Sautois et al., 2007). The parameters for the dendrite/soma and axonal compartments are identical, except that the maximum conductance values of all active channels are increased by a factor of five in the axonal compartment. The capacitance of each compartment is 5pF, and the inter-compartment conductance is 10nS. We use a two-compartment model because single compartment neurons with motoneuron properties are not able to produce persistent rhythmic firing when excited by glutamatergic synapses with NMDA receptors. Strong excitatory input leads to depolarization which stops spiking due to the depolarization block. A more realistic model incorporating two compartments is free from this problem.

### Synaptic Currents

The model neuron’s synaptic inputs *I*_*syn*_ in equation (1) includes different types of excitatory and inhibitory synapses.

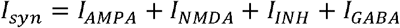

Two of these models mimic excitatory synapses by releasing glutamate and activating AMPA and NMDA receptors to the post-synaptic neurons. Two models mimic inhibitory synapses releasing glycine (INH) and Gamma-Aminobutyric acid (GABA) and activating inhibitory receptors to the post-synaptic neurons. The type of receptor released in each connection defined by the anatomical model type depends on the pre-synaptic neuron.

Each synaptic input of a post-synaptic neuron *j* is given by:

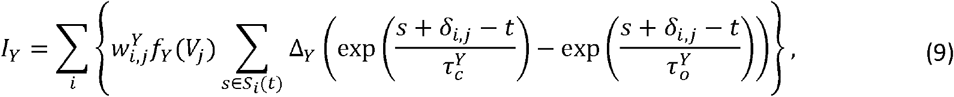

where *Y* = *AMPA, NMDA, INH, GABA*. The pre-synaptic neuronal type determines the synapse type *Y*. Glycine-releasing inhibitory synapses have presynaptic neuron types *cIN* and *aIN*; GABA-A – releasing inhibitory synapses have the presynaptic neuron type *MHR*; glutamate-releasing excitatory ones have pre-synaptic neuron types of any of the remaining cell types. Parameter 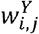 is the maximum conductance (“strength”) of synaptic connection of type Y from neuron *i* to neuron *j*. If the connectome does not include a connection from *i* to *j* then 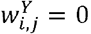, otherwise it is selected according to the type of the pre- and post-synaptic neuron types by taking into account previous experimental and modeling studies, when available. In the case that such data is not available, we explored the parameter space in order to find ranges of values that gave the desired behavior (see explanation below for each pair of connections).

*S*_*j*_ (*t*) is the set of spike times of neuron *j* up to time *t*. Each spike generates a post-synaptic current with rising time constant 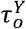 and decaying time constant 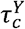. The after-spike increment Δ_Y_is a constant selected for each type of synapse except for the depressing and saturating synapses discussed below. These constants were selected so that the peak of exponentials’ difference is 1 in equation (9), meaning that following a spike the conductance rises to a maximum of 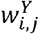. The values selected for the time constants, reverse potential and constant increments are given in Supplementary Table 3 and are based on previous models established from pairwise recordings (Hull et al., 2016; Roberts et al., 2014; Sautois et al., 2007).

### Depressing synapses

As discussed below some features of the tadpole swimming behaviour we include synaptic depression in some of the AMPA and NMDA synapses by multiplying the synaptic strength of each synapse by the depression rate at the occurrence of each pre-synaptic spike (exponentially decreasing strengths (Abbott et al., 1997)). The depressing synapses in the *CNS* model are reported in Supplementary Table 4.

### Saturating synapses

Simulations of the neural responses to trunk skin stimulation guided our construction of model synapses. To avoid the transitions from the co-activation of left-right antagonist muscles (synchrony) to the rest whilst favouring the transitions from synchrony to swimming alternations we included synaptic saturation in the *dIN* to *dIN* NMDA synapses (see Results). Indeed transitions from synchrony to rest are rare in experiments (Li et al., 2014b) and in this paper the synaptic saturation is modelled following by including a variable, rather than constant, after-spike increment Δ_NMDA_ defined by:

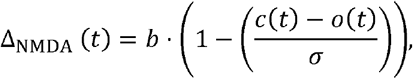

where *b* = 1.25 is the standard increment of NMDA synapses (Supplementary Table 3), σ is the level of saturation (σ=0.025), and variables 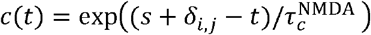 and 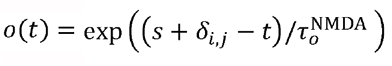 appearing in equation (9).

The function *f*_*X*_(*V*) in equation (9) describes the dependence on the post-synaptic voltage. For *Y* = *AMPA,INH* and *GABA* synapses this has a simple linear (ohmic) form:

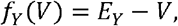

where *E*_*Y*_ is the equilibrium (reversal) potential of the synapse type. NMDA synapses have a non-linear voltage dependence term representing magnesium block and modelled by a sigmoidal scaling factor:

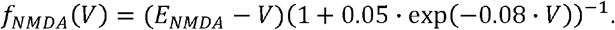

Except connections to and from *exINs* the delay terms δ_*i,j*_ in equation (9) depend on the rostro-caudal positions of the pre- and post-synaptic neurons. More, precisely, the synaptic delay between two neurons, δ_*i,j*_ consists of a constant and distance-dependent part:

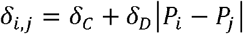

Here, *P*_*i*_ and *P*_*j*_ are the positions of neurons *i* and *j* along the rostro-caudal axis, δ_*C*_ is the constant delay and δ_*D*_ is the speed of synaptic transmission. We set δ_*C*_ = 1*ms* and δ_*D*_ =0.0035*ms*/*μm*.

We prescribe constant delays in connections to and from *exINs* because there is no available data on their rostro-caudal positions. All connections have constant delay δ_*i,j*_ = 1*ms*, except from commissural *exINs* connections, which have delay δ_*i,j*_ = 2*ms* to take into account longer distances of commissural growing axons.

### Connections types and strengths

For each connection in the adjacency matrix described in section M1 we prescribe a synaptic connection with excitatory or inhibitory active receptors depending on the pre- and post-synaptic neural types. The receptor components and their strengths were prescribed based on several experimental and/or modelling sources (see below). Supplementary Table 4 summarizes all synapses and their synaptic strength values. Depressing synapses and the citations of the papers used to infer these values are also reported in this table. To mimic synaptic strength variability, the Gaussian noise with the standard deviation 5% of the mean synaptic strength was added to the maximum conductance of all individual synapses, except for the synapses from *exIN*s and *RB*s (see below).

### Synapses from *RBs* to *dlas/dlcs*

Following (Roberts et al., 2014) we included an AMPA component and decreased the mean strength from 8nS to 4nS to distinguish between low/high level stimuli and increase the standard deviation by a factor 10 to increase the trial-to-trial variability and reduce the appearances of synchrony.

### Synapses from *dla/dlc* to *exINs*

Following (Koutsikou et al., 2018), we hypothesized that these synapses have both AMPA and NMDA components. There is no available data to infer their strengths. We thus explored the space of parameters and found a set of parameters that can explain the long and variable summation of excitation to *dIN* leading to swimming by producing (1) a long and variable firings in *exIN*s following trunk touch stimuli and (2) make swimming start more reliably on the contralateral side. The selected mean (std) strength of both AMPA and NMDA synapses from *dla* neurons is 7nS (5nS), while from *dlc* neurons is 4.2 (5nS).

### Synapses from *exINs* to *exINs*

Following the model (Koutsikou et al., 2018) we hypothesized mixed AMPA and NMDA components with modified mean strengths (from 1.5nS to 6.5nS for AMPA and from 1.8nS to and 1.4nS for NMDA) to balance the difference in the connectivity between the two modes. The strength is also scaled randomly by a chosen value (0.8, 0.6, 0.4, 0.2 or 0) to introduce trial-to-trial variability. In addition we include synaptic depression with rate 0.993 in both AMPA and NMDA components to stop the irregular firing of the *exIN*s on both sides after ∼1.5s in response to trunk skin stimuli. As a result the *hdIN* wave of depolarization generated by *exIN* EPSPs matches the wave of *hdIN* excitation in trials where swimming was not initiated (see Results and Fig. 4).

### Synapses from *exINs* to spinal *CPG* interneurons

Following (Koutsikou et al., 2018) we hypothesized mixed AMPA and NMDA components with modified mean strengths (from 0.25nS to 1.4nS for AMPA and from 0.1nS to and 0.7nS for NMDA) to balance the reduced number of connections in the model proposed here. Also these strengths are scaled randomly by a chosen value from the set (0.8, 0.6, 0.4, 0.2 or 0) to introduce trial-to-trial variability. Both AMPA and NMDA components include depression with rate 0.98, which was selected to match the *hdIN* wave of depolarization for stimuli below swimming threshold (see Results).

### Synapses from *tSts* to spinal *tINs/rdlcs*

We hypothesized that *tSt* to *rdlc* have the same mean strength as *RB* to *dlc* connections of 4nS because they are both reliable connections from sensory to sensory pathway neurons (Buhl et al., 2012; Roberts et al., 2014). For *tSt* to *tIN* connections we know from experiment that these connections include mixed AMPA and NMDA components. We include them and fit the strength of these connections to match 10-90% rise time, time to peak and duration at 50% of presumed single EPSP from whole cell recordings (Supplementary Table 5). The analysis is provided separately for only the AMPA component (obtained by NMDA block via NBOX application) and for mixed AMPA and NMDA components (Buhl et al., 2012). We fit both sets of data using a python derivative-free optimization algorithm (Mayer, 2017) that minimizes the sum of the difference between all the measures in Supplementary Table 5. The algorithm’s convergence was tested using several initial conditions. Simulated EPSPs are obtained by modeling a single connection from one *tSt* to one *tIN* and by injecting a brief step current to the *tSt* to activate a single spike. The model measures reported in Supplementary Table 4 report the values obtained by the best fitting values of AMPA and NMDA strengths and the experimental ones are in good agreement.

### Synapses from *tINs* to *hdINs*

Following pairwise whole cell recordings we know that these connections include mixed AMPA and NMDA components (Buhl et al., 2012). We fit the strength of these connections with the analysis of the 10-90% rise time and time to peak in single EPSPs from whole cell recordings. Data from these connections were only analyzed from mixed AMPA and NMDA components and from pairwise recordings of *tINs* and *dINs*. We are therefore confident that they correspond to single EPSPs from *tINs* to *dINs*. We use the same optimization technique as in *tSt* to *tIN* connections to find the best fitting strengths (see above) to match the measures of EPSPs (Supplementary Table 6). Model EPSPs are generated by assuming single synaptic connection from one *tIN* to one *dIN* and by injecting a brief step current to the *tSt* to activate a single spike. Since the *dIN* voltage dynamics depends on the electrical coupling with other *dINs* in these simulations we included 10 electrically coupled *dINs* with coupling strength 0.2nS.

### Synapses from *tINs* to *exINs*

We hypothesized that these connections are equal to *dla* to *exIN* connections, since they are both connections from sensory pathway cells to *exINs* and thus have AMPA and NMDA components with the same strengths.

### Synapses from spinal *CPG* interneurons

Synapses from *cIN*s and *aIN*s have a glycinergic INH component, while the excitatory glutamatergic synapses from *dIN*s and *mn*s have an AMPA component. The strengths of both components are based on previous models that fit whole cell recordings (Sautois et al., 2007). The AMPA strength of *dIN* to *aIN* connections was reduced (0.6nS to 0.1nS) (Roberts et al., 2014) and *aIN*s were limited to fire shortly after stimulation, as in experiments. Synapses from *dIN*s to *dIN*s include an additional NMDA component with synaptic depression with rate 0.995 to enable the spontaneous termination of swimming activity. The mean initial strength of this component is increased from 0.15nS to 0.2nS to obtain stable alternating oscillations in absence of depression.

### Synapses from *tSps* to *MHRs*

Physiological experiments (Perrins et al., 2002) provide indirect evidence that these connections are excitatory. We hypothesized that they release glutamate (as most excitatory connections in the tadpole) and include mixed AMPA and NMDA components. We selected mean connections strengths 5nS and 1nS, respectively.

### Synapses from *MHRs* to spinal neurons

Pairwise whole cell recordings revealed that these synapses release GABA-A (Perrins et al., 2002). We therefore model a GABA component and use the same mean strength (3nS) as proposed in a previous model (Hull et al., 2016).

### Statistical analysis of simulations with high and low level trunk touch and head touch

Here we report some details of *CNS* model responses to trunk touch (TT) and head touch (HT) for the proposed low and high level of stimulations across 100 repeated simulations of the model. For each protocol of stimulation we report the median for the number of activated pathway neurons; the percentage of simulations where swimming is initiated (reliability of swimming start); the median of time delays to the first reaction; and the percentage of initiations where swimming starts on the stimulated side (first flexion on stimulated side). In brackets we show the standard deviation.

- **High TT**: Stimulation of 4 *RBs*. Activated sensory pathway neurons: median is 20.5 *dlas* (2.8) and 46 *dlcs* (6.7). Reliability of swimming start: 78%. Delay: median is 52.4 ms (13.4 ms). First flexion on stimulated side: 37%.
- **Low TT**: Stimulation of 2 *RBs*. Activated sensory pathway neurons: median is 11 *dlas* and 23 *dlcs*. Reliability of swimming start: 52%. Delay: median is 68.7 ms (18.7 ms). First flexion on stimulated side: 37%.
- **High HT**: Stimulation of 4 *tSts*. Activated sensory pathway neurons: median is 31 *dlcs* (2.2) and 15 *tINs* (2.2). Reliability of swimming start: 95%. Delay: median is 17.7 ms (1.9 ms). First flexion on stimulated side: 100%.
- **Low HT**: Stimulation of 2 *tSts*. Activated sensory pathway neurons: median is 21 *dlcs* (3.2) and 6 *tINs* (2.3). Reliability of swimming start: 68%. Delay: median is 57.7 ms (24.5 ms). First flexion on stimulated side: 67%.

### Transformation of motoneuron spiking times to muscle contraction forces

Each motoneuron’s single spike which occurs at time *t*_*spike*_ (ms) causes each ELM (elementary muscle fiber) within innervated muscle segment to contract with the force

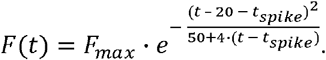

Where *F*_*max*_ was chosen such that it provides the most realistic movement.

Summation of forces. Assume:

- One spike appears at *t*_*prev_spike*_ and it is followed by another spike at *t*_*spike*_,
- The resting activity of the muscle segment at the beginning of the second spike at time *t*_*spike*_ is 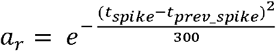.

Then the force caused by these two spikes is calculated as

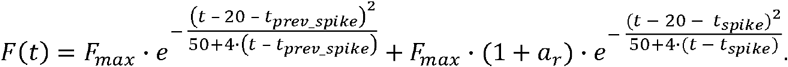

In case of more spikes the impact from each new spike is calculated in the same way.

## Acknowledgments

We are grateful to Robert Merrison-Hort and Edgar Buhl for valuable feedback on the models’ developments. **A.F**. acknowledges support from the UK Engineering and Physical Sciences Research Council (EPSRC) New Investigator Award (EP/R03124X/1). **A.P**. acknowledges support from the Russian Federation state assignment IIS SB RAS. **S.K**. acknowledges support from the UK Physiological Society. **R.B**. acknowledges support from the UK Biotechnology and Biological Sciences Research Council (BBSRC): BB/L000814/1, BB/T002352/1. **W.L**. acknowledges support from the UK Biotechnology and Biological Sciences Research Council (BBSRC): BB/T003146.

## Author Contributions

Conceptualization: A.F., A.P., A.R., S.S., and R.B.; software development and simulations: A.F. (CNS model) and A.P. (VT model); formal analysis: A.F. and A.P.; supervision: R.B. and A.R.; experimental data collection: S.K., S.S., W.L.; writing: A.F., A.P., A.R., and R.B. (original draft) and A.F., A.P., R.B., A.R., W.L., and S.K. (review and editing); funding acquisition for the project: S.S, A.R., W.L., and R.B.

## Supplementary Materials

**Supplementary video1**: Swimming of 3D biomechanical Virtual Tadpole model of a stage 37/38 *Xenopus* tadpole stimulated on the left trunk at 50 ms. Body movement in the water tank seen from top and right side views. Muscle activation levels on each side are shown by red colour and numbers at top information panel. The tadpole stops when it hits the tank wall. Speed of water particles shown by colours from light green (slow) to red (fast).

**Supplementary video2**: Body rotation during swimming of 3D biomechanical Virtual Tadpole model of a stage 37/38 Xenopus tadpole. The tadpole initially lies on its right side on the tank bottom. Skin stimulation on left trunk at 50 ms leads to muscle contraction on the right side and, as swimming continues, the body rotates to a standard dorsal-up, belly-down position. Body movement in the water tank seen from top and right side views. Muscle activation levels on each side are shown by red colour and numbers at top information panel. The tadpole stops when it hits the tank wall. Speed of water particles shown by colours from light green (slow) to red (fast).

**Supplementary Table 1**: Capacitance, maximal conductance and equilibrium potential of each ionic channel in the model neurons.

**Supplementary Table 2**: Parameters of the rate functions for the Hodgkin – Huxley type neuronal model for all neuron types.

**Supplementary Table 3**: Parameters of the synaptic models for NMDA, AMPA and INH synapses.

**Supplementary Table 4**: Summary of all the synaptic connections between different neuronal populations.

**Supplementary Table 5**: Measures of *tSt* to *tIN* EPSPs in model simulations and experiment.

**Supplementary Table 6**: Measures of single *tIN* to *dIN* EPSPs in model simulations and experiment.

