## Supplementary Table 1 for "Whole animal modelling reveals neuronal mechanisms of decision-making and reproduces unpredictable swimming in frog tadpoles"

|  | $\mathbf{C}$ (pF) | $\boldsymbol{g}_{\boldsymbol{lk}}$ (nS) | $\boldsymbol{e}_{\boldsymbol{lk}}\mathbf{(mV)}$ | $\boldsymbol{g}_{\boldsymbol{Na}}\mathbf{(nS)}$ | $\boldsymbol{e}_{\boldsymbol{Na}} \mathbf{(mV)}$ | $\boldsymbol{g}_{\boldsymbol{Kf}}\mathbf{(nS)}$ | $\boldsymbol{e}_{\boldsymbol{Kf}}\mathbf{(mV)}$ | $\boldsymbol{g}_{\boldsymbol{Ks}}\mathbf{(nS)}$ | $\boldsymbol{e}_{\boldsymbol{Ks}}\mathbf{(mV)}$ |
| --- | --- | --- | --- | --- | --- | --- | --- | --- | --- |
| dIN | 10 | $2.5$ | $-52$ | $300$ | $50$ | $25$ | $-81.5$ | $20$ | $-81.5$ |
| dla/dlc/aIN/cIN/mn | 10 | $2.5$ | $-61$ | $110$ | $50$ | $8$ | $-80$ | $1$ | $-80$ |
| RB/tSt/tSp | 4 | $4.4$ | $-60$ | $120$ | $50$ | $1.5$ | $-80$ | $8$ | $-80$ |
| tIN | 4 | $2.2$ | $-55$ | $680$ | $50$ | $40$ | $-80$ | $20$ | $-80$ |
| MHR | 4 | $3.8$ | $-60$ | $420$ | $50$ | $70$ | $-80$ | $10$ | $-80$ |

Supplementary Table 1. Capacitance, maximal conductance and equilibrium potential of each ionic channel in the model neurons.
