## Supplementary Table 2 for "Whole animal modelling reveals neuronal mechanisms of decision-making and reproduces unpredictable swimming in frog tadpoles"

| dIN | Rate Function | | | | A ($\boldsymbol{m}\boldsymbol{s}^{\boldsymbol{-1}}$) | | | B ($\boldsymbol{m}\boldsymbol{s}^{\boldsymbol{-1}}\boldsymbol{m}\boldsymbol{V}^{\boldsymbol{-1}}$) | | | C (-) | | | D ($\boldsymbol{mV}$) | | | E ($\boldsymbol{mV}$) |
| --- | --- | --- | --- | --- | --- | --- | --- | --- | --- | --- | --- | --- | --- | --- | --- | --- | --- |
| Ca | $\alpha_{r}$ | | | | $4$ | | | $0$ | | | $1$ | | | $-15.3$ | | | $-13.6$ |
|  | $\beta_{r}(v<-25mV)$ | | | | $1.2$ | | | $0$ | | | $1$ | | | $10.6$ | | | $1$ |
|  | $\beta_{r}(v>-25mV)$ | | | | $1.3$ | | | $0$ | | | $1$ | | | $5.4$ | | | $12.1$ |
| K-fast | $\alpha_{f}$ | | | | $5.1$ | | | $0.1$ | | | $5.1$ | | | $-18.4$ | | | $-25.4$ |
|  | $\beta_{f}$ | | | | $0.5$ | | | $0$ | | | $0$ | | | $28.7$ | | | $34.6$ |
| K-slow | $\alpha_{s}$ | | | | $0.5$ | | | $8.2e-3$ | | | $4.6$ | | | $-4.2$ | | | $-12$ |
|  | $\beta_{s}$ | | | | $0.1$ | | | $-1.3e-3$ | | | $1.6$ | | | $2.1e5$ | | | $3.3e5$ |
| Na | $\alpha_{m}$ | | | | $8.7$ | | | $0$ | | | $1$ | | | $-1$ | | | $12.6$ |
|  | $\beta_{m}$ | | | | $3.8$ | | | $0$ | | | $1$ | | | $9$ | | | $9.7$ |
|  | $\alpha_{h}$ | | | | $0.1$ | | | $0$ | | | $0$ | | | $38.9$ | | | $26$ |
|  | $\beta_{h}$ | | | | $4.1$ | | | $0$ | | | $1$ | | | $-5.1$ | | | $-10.2$ |
| dla/dlc/aIN/  cIN/mn | | Rate Function | A ($ms^{-1}$) | | | B ($ms^{-1}mV^{-1}$) | | | C (-) | | | D ($mV$) | | | E ($mV$) | | |
| K-fast | $\alpha_{f}$ | | | | $3.1$ | | | $0$ | | | $1$ | | | $-27.5$ | | | $-9.3$ |
|  | $\beta_{f}$ | | | | $0.4$ | | | $0$ | | | $1$ | | | $9$ | | | $16.2$ |
| K-slow | $\alpha_{s}$ | | | | $0.2$ | | | $0$ | | | $1$ | | | $-3$ | | | $-7.7$ |
|  | $\beta_{s}$ | | | | $0.05$ | | | $0$ | | | $1$ | | | $-14.1$ | | | $6.1$ |
| Na | $\alpha_{m}$ | | | | $13.3$ | | | $0$ | | | $0.5$ | | | $-5.$ | | | $-12.6$ |
|  | $\beta_{m}$ | | | | $5.7$ | | | $0$ | | | $1$ | | | $5$ | | | $9.7$ |
|  | $\alpha_{h}$ | | | | $0.04$ | | | $0$ | | | $0$ | | | $28.8$ | | | $26$ |
|  | $\beta_{h}$ | | | | $2$ | | | $0$ | | | $0$ | | | $-9.1$ | | | $-10.2$ |
| RB/tSt/tSp | Rate Function | | | A ($ms^{-1}$) | | | B ($ms^{-1}mV^{-1}$) | | | C (-) | | | D ($mV$) | | | E ($mV$) | |
| K-fast | $\alpha_{f}$ | | | | $3.1$ | | | $0$ | | | $1$ | | | $-32.5$ | | | $-9.3$ |
|  | $\beta_{f}$ | | | | $0.4$ | | | $0$ | | | $1$ | | | $4$ | | | $16.2$ |
| K-slow | $\alpha_{s}$ | | | | $0.2$ | | | $0$ | | | $1$ | | | $-8$ | | | $-7.7$ |
|  | $\beta_{s}$ | | | | $0.05$ | | | $0$ | | | $2$ | | | $-19.1$ | | | $6.1$ |
| Na | $\alpha_{m}$ | | | | $13.01$ | | | $0$ | | | $1$ | | | $-1$ | | | $-12.6$ |
|  | $\beta_{m}$ | | | | $5.7$ | | | $0$ | | | $1$ | | | $6$ | | | $9.7$ |
|  | $\alpha_{h}$ | | | | $0.06$ | | | $0$ | | | $0$ | | | $30$ | | | $26$ |
|  | $\beta_{h}$ | | | | $2$ | | | $0$ | | | $1$ | | | $-8.1$ | | | $-10.2$ |
| tIN | Rate Function | | | A ($ms^{-1}$) | | | B ($ms^{-1}mV^{-1}$) | | | C (-) | | | D ($mV$) | | | E ($mV$) | |
| K-fast | $\alpha_{f}$ | | | | $3.1$ | | | $0$ | | | $1$ | | | $-50.5$ | | | $-9.3$ |
|  | $\beta_{f}$ | | | | $1.1$ | | | $0$ | | | $1$ | | | $-16$ | | | $16.2$ |
| K-slow | $\alpha_{s}$ | | | | $0.2$ | | | $0$ | | | $1$ | | | $-26$ | | | $-7.7$ |
|  | $\beta_{s}$ | | | | $0.05$ | | | $0$ | | | $1$ | | | $-37.1$ | | | $6.1$ |
| Na | $\alpha_{m}$ | | | | $8.67$ | | | $0$ | | | $0.5$ | | | $-28$ | | | $-18.6$ |
|  | $\beta_{m}$ | | | | $3.7$ | | | $0$ | | | $1$ | | | $-12$ | | | $9.7$ |
|  | $\alpha_{h}$ | | | | $0.04$ | | | $0$ | | | $0$ | | | $0.8$ | | | $26$ |
|  | $\beta_{h}$ | | | | $4.1$ | | | $0$ | | | $0$ | | | $-35$ | | | $-10.2$ |
| MHR | Rate Function | | | A ($\boldsymbol{m}\boldsymbol{s}^{\boldsymbol{-1}}$) | | | B ($\boldsymbol{m}\boldsymbol{s}^{\boldsymbol{-1}}\boldsymbol{m}\boldsymbol{V}^{\boldsymbol{-1}}$) | | | C (-) | | | D ($\boldsymbol{mV}$) | | | E ($\boldsymbol{mV}$) | |
| K-fast | $\alpha_{f}$ | | | | $3.1$ | | | $0$ | | | $1$ | | | $-42.5$ | | | $-9.3$ |
|  | $\beta_{f}$ | | | | $0.2$ | | | $0$ | | | $2$ | | | $-6$ | | | $16.2$ |
| K-slow | $\alpha_{s}$ | | | | $4$ | | | $0$ | | | $1$ | | | $-63$ | | | $-7.7$ |
|  | $\beta_{s}$ | | | | $0.01$ | | | $0$ | | | $1$ | | | $37$ | | | $6.1$ |
| Na | $\alpha_{m}$ | | | | $16.3$ | | | $0$ | | | $3$ | | | $-13$ | | | $-12.6$ |
|  | $\beta_{m}$ | | | | $5.7$ | | | $0$ | | | $1$ | | | $-4$ | | | $9.7$ |
|  | $\alpha_{h}$ | | | | $0.06$ | | | $0$ | | | $0$ | | | $9.9$ | | | $26$ |
|  | $\beta_{h}$ | | | | $4.1$ | | | $0$ | | | $0$ | | | $-18.1$ | | | $-10.2$ |

Supplementary Table 2: Parameters of the rate functions for the Hodgkin – Huxley type neuronal model for all neuron types. Parameter values are rounded to the first decimal digit.
