## Supplementary Table 3 for "Whole animal modelling reveals neuronal mechanisms of decision-making and reproduces unpredictable swimming in frog tadpoles"

| $\boldsymbol{X}$ | **NMDA** | **AMPA** | **INH** | **GABA** |
| --- | --- | --- | --- | --- |
| $\boldsymbol{\tau}_{\boldsymbol{o}}^{\boldsymbol{X}}\mathbf{(}\boldsymbol{ms}\mathbf{)}$ | $0.5$ | $0.2$ | $1.5$ | $1.5$ |
| $\boldsymbol{\tau}_{\boldsymbol{c}}^{\boldsymbol{X}}\mathbf{(}\boldsymbol{ms}\mathbf{)}$ | $80$ | $3$ | $4$ | $70$ |
| $\boldsymbol{\Delta}_{\boldsymbol{X}}\mathbf{(-)}$ | $1.25$ | $1.25$ | $3$ | $3$ |
| $\boldsymbol{E}_{\boldsymbol{X}}\mathbf{(}\boldsymbol{mV}\mathbf{)}$ | $0$ | $0$ | $-75$ | $-70$ |

Supplementary Table 3: Parameters of the synaptic models for NMDA, AMPA and INH synapses.
