## Supplementary Table 4 for "Whole animal modelling reveals neuronal mechanisms of decision-making and reproduces unpredictable swimming in frog tadpoles"

| Connection | Components | <Value/s (nS)> | Depression | Source |
| --- | --- | --- | --- | --- |
| *RB->dla/dlc* | AMPA | 4 | No | Roberts et al., 2014 |
| *dla ->exIN* | AMPA/NMDA | 7*/7* | No | Koutsikou et al., 2018 |
| *dlc ->exIN* | AMPA/NMDA | 4.2*/4.2* | No | Koutsikou et al., 2018 |
| *exIN->exIN* | AMPA/NMDA | 6.5*/1.4* | Yes | Koutsikou et al., 2018 |
| *exIN->CPG* | AMPA/NMDA | 1.4*/0.7* | Yes | Koutsikou et al., 2018 |
| *tSt->tIN* | AMPA/NMDA | 1/0.7 | No | Buhl et al., 2012 |
| *tSt->rdlc* | AMPA | 8 | No | Roberts et al., 2014 |
| *tIN->exIN* | AMPA/NMDA | 7*/7* | No | Koutsikou et al., 2018 |
| *tIN->hdIN* | AMPA/NMDA | 0.35/0.3 | No | Buhl et al., 2012 |
| *cIN/aIN->CPG* | INH | 0.4 | No | Roberts et al., 2014 |
| *dIN/mn->CPG* | AMPA | 0.6 | No | Roberts et al., 2014 |
| *dIN->aIN* | AMPA | 0.1 | No | Roberts et al., 2014 |
| *dIN->dIN* | NMDA | 0.2* | Yes | Roberts et al., 2014 |
| *tSp->MHR* | AMPA/NMDA | 5/1 | No | Perrins et al., 2002 |
| *MHR->CPG* | GABA | 4 | No | Hull et al., 2016 |

Supplementary Table 4: Summary of all the synaptic connections between different neuronal populations.

In order of columns we report: (1) directed connection between a pair of populations; (2) active receptor types; (3) the mean strength of each synaptic connection; (* is used to indicate that the values of connection strength was modified from the value proposed by the corresponding reference material); (4) if the synaptic connection was depressed; (5) the reference to experimental and/or modeling supporting materials for modelling the synaptic strength.
