## Supplementary Table 5 for "Whole animal modelling reveals neuronal mechanisms of decision-making and reproduces unpredictable swimming in frog tadpoles"

| tSt to tIN EPSPs | | | | | |
| --- | --- | --- | --- | --- | --- |
| NMDA | model | exp | AMPA+NMDA | model | exp |
| Maximal amplitude (mV) | 4.7 | 4.7 |  | 14.4 | 14.2 |
| 10-90% rise time (ms) | 10.1 | 10.4 |  | 1.9 | 2.5 |
| Duration at 50% amplitude (ms) | 64 | 54.4 |  | 11.21 | 17.8 |
| Time to peak (ms) | 24.8 | 16.6 |  | 10.2 | 8.7 |

Supplementary Table 5: Measures of *tSt* to *tIN* EPSPs in model simulations and experiment.

Experimental measures (exp) have been reported in Buhl et al., 2012. We show measures for NMDA component (left: model and exp) and mixed AMPA and NMDA (right: model and exp). These measures are: the average of the maximal EPSP amplitude, the rise time from 10% to 90% of the EPSP amplitude, the EPSP duration at 50% amplitude, and time to peak from the stimulus. The experimental values were analyzed in *tIN* single whole cell recordings by application of weak head stimuli that are presumed to activate a single EPSP from a *tSt*. In model simulations the strengths of the AMPA and NMDA are $w_{AMPA}=1nS$ and $w_{NMDA}=0.7nS$, respectively.
