## Supplementary Table 6 for "Whole animal modelling reveals neuronal mechanisms of decision-making and reproduces unpredictable swimming in frog tadpoles"

| tIN to dIN EPSPs | | |
| --- | --- | --- |
| AMPA+NMDA | model | exp |
| Amplitude (mV) | 2.2 | 2.6 |
| Duration at 50% amplitude (ms) | 13.8 | 14 |
| Time to peak (ms) | 3.4 | 5.2 |

Supplementary Table 6: Measures of single *tIN* to *dIN* EPSPs in model simulations and experiment. Experimental measures (exp) have been reported in Buhl et al., 2012. We show measures a mixture of AMPA and NMDA components. These measures are: the average of the maximal EPSP amplitude, the EPSP duration at 50% amplitude, and time to peak from the stimulus. The experimental values were analyzed in pairwise (*tIN,dIN*) recordings. Strengths of the model AMPA and NMDA synapses are $w_{AMPA}=0.35nS$ and $w_{NMDA}=0.3nS$, respectively. The post-synaptic *dIN*s are electrically coupled to other 10 *dIN*s with electrical coupling 0.2nS.
